# Decoding the Spatial and Epigenetic Logic of Early Skin Development

**DOI:** 10.1101/2025.09.19.677295

**Authors:** Nigel L. Hammond, Syed Murtuza Baker, Amanda McGovern, Antony Adamson, Andrew D Sharrocks, Magnus Rattray, Jill Dixon, Michael J. Dixon

## Abstract

The embryonic epidermis originates from a simple single-layered ectoderm that undergoes precise transcriptional and morphological transitions to generate a self-replenishing, multi-layered epidermis. A key early milestone is the emergence of the periderm; a transient, protective layer whose disruption underlies a series of devastating human congenital disorders. Despite its importance, the gene regulatory networks that drive periderm formation remain poorly defined.

Here, we integrate single-cell multiome profiling with high-resolution spatial transcriptomics analysis in wild-type and transcription factor p63 mutant (*Trp63*^-/-^) embryos to dissect the transcriptional and epigenetic programs driving epidermal lineage diversification. We resolve the cellular repertoire of early embryonic skin, define epithelial subpopulations and reconstruct the transcriptional dynamics underlying the periderm emergence. We show that p63 sits at the core of a multilayered enhancer-gene regulatory network, demonstrate how this network is perturbed *in vivo*, and functionally validate key targets using comparative cellular models. Together, these findings delineate the gene regulatory architecture of early skin development and establish a framework for understanding epithelial-associated congenital disorders.

## Introduction

During embryogenesis, the ectoderm transitions through a series of defined cell states to produce a self-replenishing, multi-layered epidermis that provides protection against dehydration, mechanical trauma, and microbial invasion^1^. The first stratification event occurs when ectodermal cells detach from the basement membrane to form a superficial layer of flattened periderm cells^2^.

On mouse embryonic day (E)9, the ΔNp63α-positive ectoderm expresses keratins 8/18 which are characteristic of a simple epithelium^1^. At this stage, the plane of cell division is parallel to the basement membrane and so unlike later stratification which results from asymmetric cell division^3^, periderm forms as basal cells extend cellular processes over their neighbours, detach from the basement membrane, and migrate superficially to form a surface layer of flattened cells^4,5^. Periderm formation is patterned starting on the tail and limbs before spreading over the torso and face to cover the embryo by E14^6^. All stages of periderm development are present in E10, E11 and E12 embryos. During E16-E17, periderm disaggregates due to loss of contact with the terminally differentiated keratinocytes in a pattern mirroring barrier formation^6^.

Prior to periderm formation ectodermal cells are highly polarised with adjacent cells joined by adhesion complexes, expression of which is limited by tight junctions; as a result, the cells are unable to adhere to adjacent ectodermal surfaces^6^. Subsequently, although the basal cells are no longer polarised, tight junctions on the apico-lateral surfaces of the newly formed periderm cells provide a ‘fence function’ to prevent spread of adhesion complexes onto their apical surfaces. In contrast, in the absence of periderm cell adhesion molecules are expressed on the apical surface of the exposed cells which allows inter-epithelial adhesion; thus, periderm forms a barrier to prevent pathological adhesion between adjacent epithelia^6^.

Failure of periderm formation underlies human congenital disorders characterised by webbing across the major joints, cleft lip and/or palate, digital fusions, and genital malformations^2^. These conditions include: Van der Woude, popliteal pterygium, Bartsocas Papas, and cocoon syndromes which arise from mutations in the genes encoding the transcription factors IRF6 (Van der Woude syndrome, VWS; popliteal pterygium syndrome)^7^ and GRHL3 (subset of VWS)^8^, atypical protein kinase PRKCI (subset of VWS)^9^, the receptor interacting kinase RIPK4 (Bartsocas Papas syndrome, BPS)^10,11^, and the kinase IKKα (cocoon syndrome; BPS)^12,13^. However, there are major gaps in our understanding of the gene regulatory networks driving periderm development^2^.

Here, we present a spatially resolved single cell multiome atlas of early embryonic skin development, capturing the emergence of periderm. By integrating single nucleus transcriptome (snRNA-seq) and chromatin accessibility (snATAC-seq) profiling with high-resolution spatial transcriptomics (MERSCOPE), we define the transcriptional and regulatory programs underlying epithelial differentiation and periderm formation. Through integrated pseudo-temporal modelling, we reconstruct the transcriptional dynamics of the basal to periderm transition, uncovering a hierarchical transcription factor cascade orchestrated by P63. Focussing on this master regulator, we infer the P63 enhancer-gene regulatory network, refine it through *in vivo* perturbation analyses, and functionally validate key targets using CRISPR interference in comparative cellular models. Together, these findings provide new insight into the spatial gene regulatory architecture of early skin development and establish a framework for understanding epithelial-associated congenital disorders.

## Results

### Generation of a spatially resolved single cell multiome atlas of embryonic skin development

To define the cellular and molecular processes underlying early embryonic skin development, we performed single cell profiling of the transcriptome (snRNA-seq; 3’) and accessible chromatin (snATAC-seq) from the same nucleus (sc-multiome analysis), of lateral skin dissected from wild-type mouse embryos at E10, E11 and E12 (Fig. 1A). We dissociated the tissue to single cells, and nuclei with >90% viability were isolated to ensure high-quality input for downstream processing. Using the 10X Chromium platform, we generated joint snRNA-seq and snATAC-seq libraries for each developmental age.

**Fig. 1.**
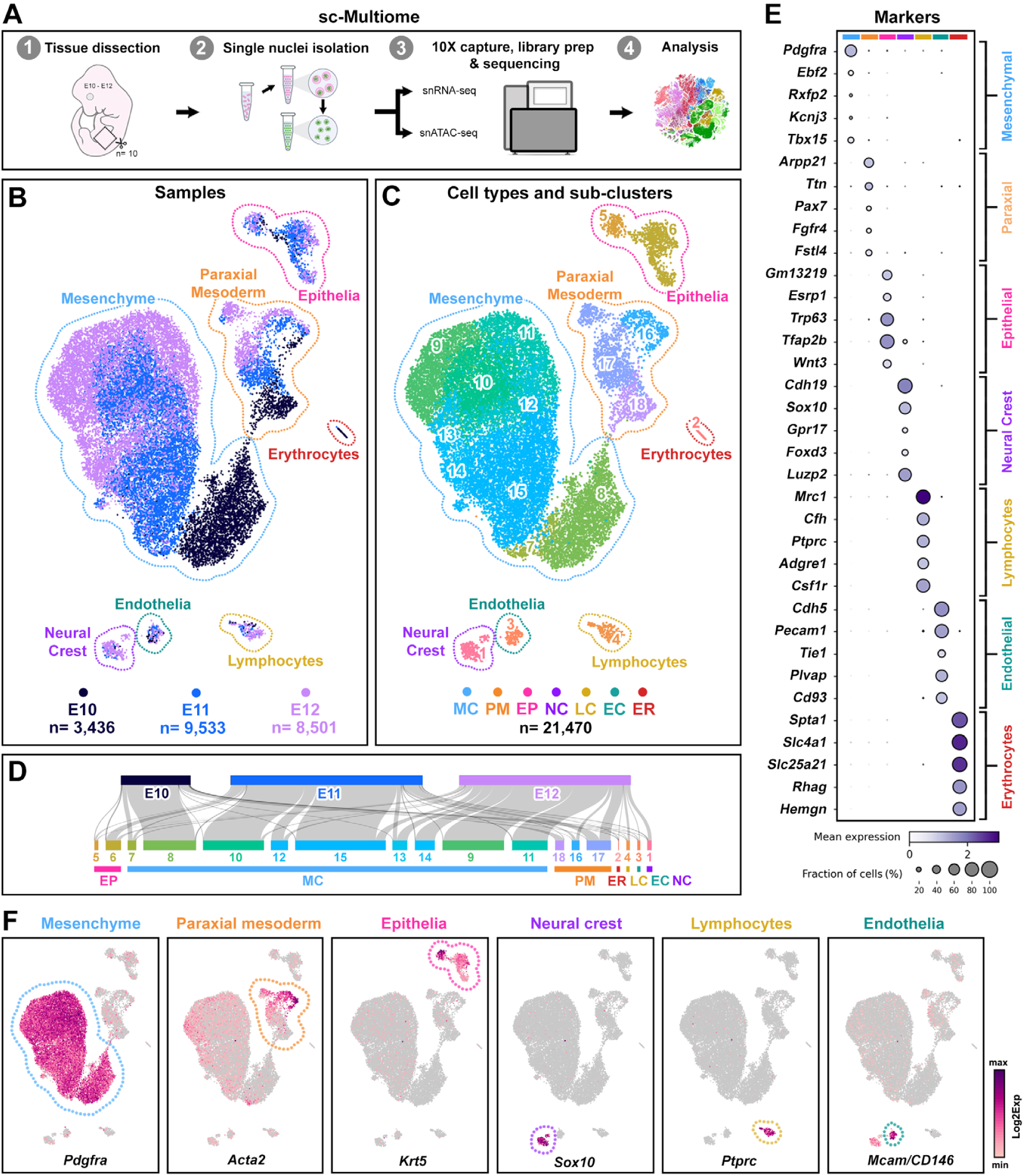
Analysis of early embryonic skin development at single cell resolution. (**A**) Generation of multiome datasets from dorsal skin dissected from E10-E12 wild-type embryos. (**B,C**) Joint clustering of multimodal data (E10-E12) from 21,470 cells reveals seven major cell types and 18 cell sub-populations. (**D**) Cell proportions for samples, sub-clusters and cell types. (**E,F**) Dot plots and expression UMAPs of cell type marker genes. EP epithelia, MC mesenchyme, PM paraxial mesoderm, EC endothelia, NC neural crest, LC lymphocytes, ER erythrocytes.

Following sequencing, read mapping, and dataset aggregation, we performed quality control, joint modality (sub)clustering, marker gene analysis, and cell type annotation to characterise the cell populations. Using gene expression profiling from 21,470 nuclei, we captured an average of 3,497 genes and 7,193 UMIs per cell (Fig. 1; Extended Data Fig. 1A-E). Profiling snRNA-seq data, we defined marker genes per cell cluster, and annotated seven major cell types: epithelia (*Trp63, Krt14, Krt5, Gm13219*), mesenchyme (*Pdgfra*, *Pdgfrb, Col24a1, Tbx15*), paraxial mesoderm (*Ttn, Fgf9, Pax3, Acta2*), lymphocytes (*Mrc1*, *Ptprc*, *Csf1r, F13a1*), endothelia (*Cdh5, Plvap, Tie1, Mcam*), neural crest (*Cdh19*, *Sox10, Foxd3, Gpr17*) and erythrocytes (*Spta1*, *Slc4a1*, *Rhag, Sptb*).

Using Leiden clustering, these broad cell types further resolved to 18 transcriptionally distinct sub-populations, which when considered with developmental time-points, broadly revealed the trajectories of these lineages (Fig. 1B-D; Extended Data Fig 1A,B; Supplementary Table 1). For example, mesenchymal sub-populations encompassed early (E10; clusters 7,8), mid (E11; clusters 10,12,15) and late (E12; clusters 9,11) development. Similarly, clusters 16,17 and 18 (E10-E12) reflected development of the paraxial mesoderm (Fig. 1B,D). Together, our dataset captured both abundant cells, such as mesenchymal fibroblasts (83%), and rare sub-populations, such as neural crest-derived Schwann cell precursors^14^ (*Sox10*, *Dlx1*, *Dlx2*; 0.9%), melanocytes (*Pmel, Dct, Tyr;* 0.1%) and epithelial periderm (*Gabrp*, *Nectin4, Cldn23*; 0.9%) (Extended Data Fig. 1E).

While single cell sequencing enabled high-resolution profiling of cellular diversity, the spatial organisation of these populations within the developing skin was lost due to tissue dissociation. To address this issue, we employed spatial transcriptomics which allows for gene expression profiling while preserving tissue architecture. We used the MERSCOPE platform to perform *in situ* spatial imaging of transverse mid-trunk sections from E12 embryos at sub-cellular resolution. A custom 285-gene panel was designed, incorporating up to 20 marker genes per cell type identified in the single-cell atlas, along with additional genes relevant to epidermal development curated from the literature, ensuring the greatest power to spatially delineate cell sub-types (Supplementary Table 2).

Subsequently, we generated three spatial transcriptomics datasets (samples 1-3; Extended Data Fig. 2) capturing regions corresponding to those profiled in the sc-multiome analysis. Individual RNA molecules were visualised across seven z-stack focal planes and assigned to cells segmented using a combination of three cell membrane protein stains, DAPI, and total RNA signals. This approach enabled the reconstruction of single-cell expression profiles, with ∼3.5 million transcripts detected in 57,799 segmented cells.

Using sample 2 as a representative dataset, we applied quality control and Leiden clustering to identify 28 spatial clusters among 18,942 cells (Fig. 2A). By correlating spatial clusters with marker gene profiles from our sc-multiome atlas, we revealed the spatial distribution of distinct cell populations (Fig. 2A-F; Extended Data Fig. 3, 4). For example, neural crest-derived spatial cluster 21 was enriched for melanocyte markers (*Pmel, Sox10*) and localised near the epidermis, whereas cluster 27 corresponded to more abundant populations of Schwann cell precursors (*Sox10*, *Dlx2*) dispersed throughout the dermis (Fig. 2E). The broader tissue coverage of spatial imaging allowed us to identify eight ‘*de novo*’ clusters, which we annotated based on shared marker expression and spatial localisation, such as midgut epithelium (clusters 9, 10, 25; *Krt7*, *Krt8*, *Epcam*), neural tube populations (cluster 26; *Pax3*), and chondroprogenitors (cluster 12; *Sox9, Sox5*) (Fig. 2F).

**Fig. 2.**
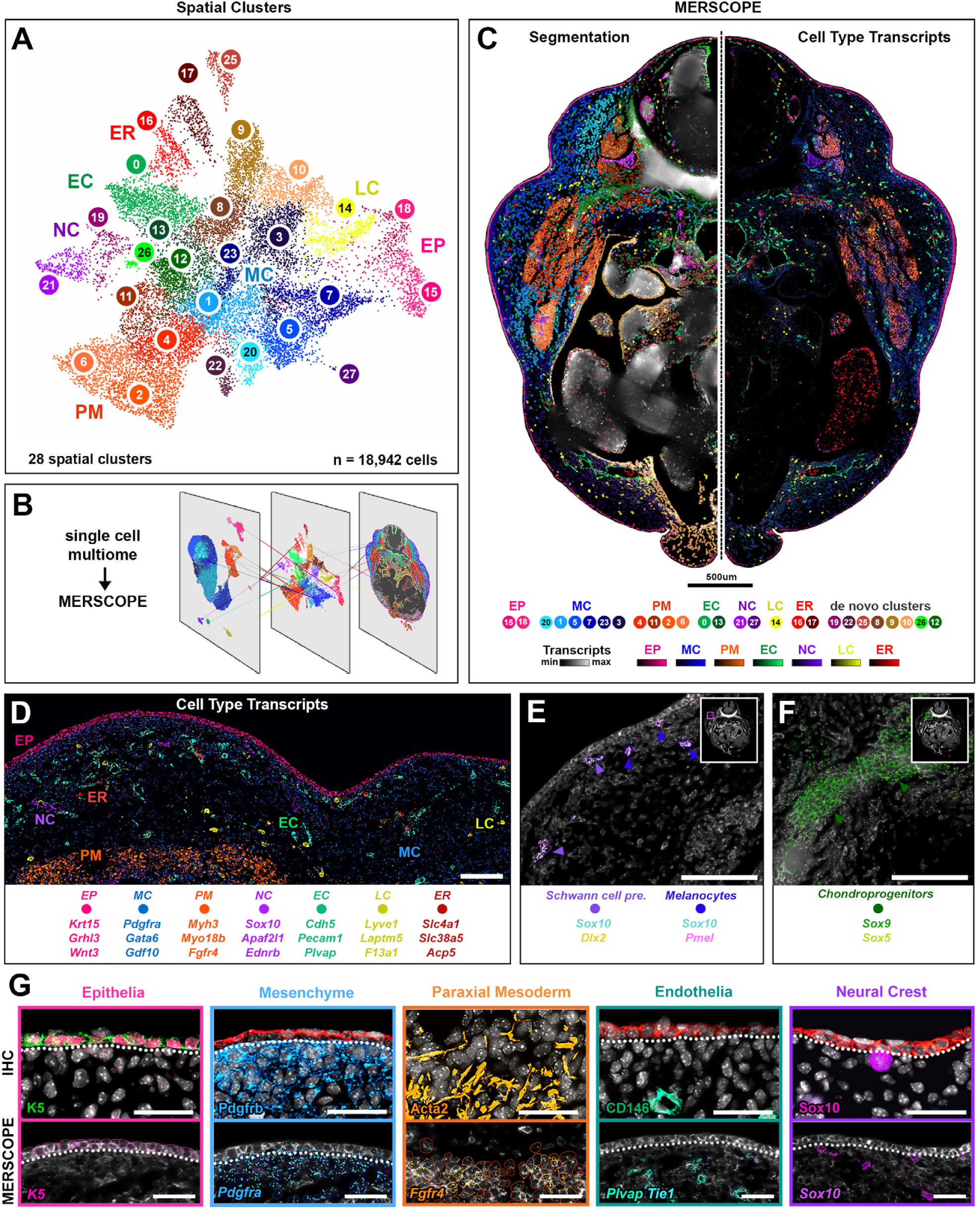
Integration of spatial transcriptomics and sc-multiome data. (**A-F**) Spatial *in situ* imaging (MERSCOPE) of transverse tissue sections (E12) showing transcripts, clustering and segmentation of single cells. (**A,B**) UMAP spatial clustering of segmented cells (sample 2) and integration with sc-multiome. (**C**) Transverse tissue section (mirrored along dotted line) showing cell segmentation of spatial clusters, coloured by cell sub-type, with average transcript density of marker genes per cell type. (**D**) Higher resolution of the region profiled for sc-multiome, showing transcripts of several genes that are cell type-specific. (**E,F**) Spatial mapping of neural crest subpopulations (**E**) and chondroprogenitors (**F**). (**G**) Spatial expression correlation between protein and transcripts that are cell type-specific. Scalebars D-F, 100 µm; G, 50 µm

To validate our integrated spatially resolved single-cell atlas, we performed immunofluorescence analysis for selected markers, confirming that protein expression patterns correlated with spatial transcriptomic profiles of cell type marker genes (Fig. 2G).

### Integrative single-cell multiome analyses reveal cell lineage-defining TFs

To investigate the regulatory landscape of embryonic skin development, we examined chromatin accessibility using snATAC-seq data, analysed through joint modality clustering (Extended Data Fig. 1A,B). Across all developmental stages, we annotated accessible regions (peaks) to subclusters of the major cell types and identified regions of open chromatin that were preferentially enriched near marker genes of each lineage (Fig. 3A).

**Fig. 3.**
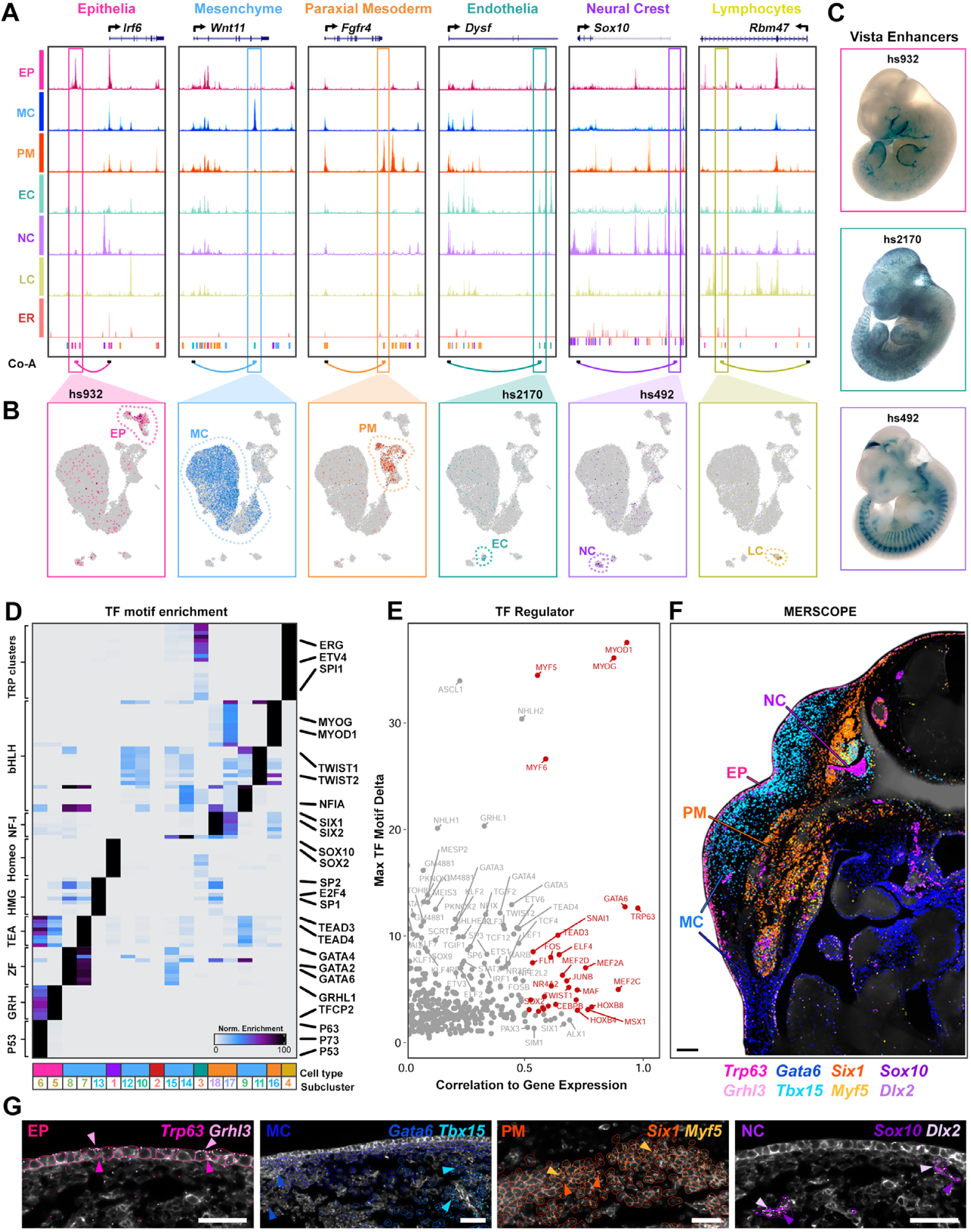
Joint modality single cell analyses reveal cell lineage defining transcription factors. (**A,B**) Cell type snATAC-seq signal tracks of marker gene loci reveal enhancers that are co-accessible (co-A) with promoters. UMAP plots of cut-site counts demonstrate cell specificity. (**C**) Transient transgenic assays (VISTA enhancers) of cell-type specific enhancers showing wholemount expression of *lacZ* in epithelia (hs932), endothelia (hs2170) and neural crest-derived somites (hs492). (**D**) Transcription factor (TF) motif enrichment on cell (sub)type snATAC-seq peaks. (**E**) Volcano plot showing correlation between enriched motifs and gene expression across all cells. (**F,G**) Spatial transcriptomics (MERSCOPE) on E12 transverse tissue section, showing spatial organisation of transcripts for lineage defining TFs. *Trp63/Grhl3*, epithelia; *Gata6/Tbx15*, mesenchyme; *Six1/Myf5*, paraxial mesoderm; *Sox10/Dlx2*, neural crest. Scalebars F,G 50 µm.

Leveraging joint transcript and open chromatin measurements from the same cell, we used co-accessibility inference (Cell Ranger ARC) to predict putative enhancer-promoter interactions and assess their correlation with gene expression. This analysis yielded 121,153 significant peak-to-gene linkages (correlation >0, -log10fdr >5), enabling the identification of cell type-specific regulatory elements (Supplementary Table 3). For example, we detected a well-characterised disease-associated epithelial-specific enhancer located ∼12 kb from the *Irf6* locus and co-accessible with its promoter. This element (Vista enhancer hs932), and disease-causing variants within, have been validated as *bona fide* enhancers *in vivo* using transient transgenic reporter assays (Fig. 3A-C)^15–17^. Similarly, we identified novel enhancer-gene linkages for additional lineage-specific loci, including an endothelial enhancer (Vista enhancer hs2170) linked to *Dysf* expression in developing vasculature^18^, and a neural crest-specific enhancer (Vista enhancer hs492) associated with *Sox10* expression in populations of Schwann cell precursors (Fig. 3A-C)^14^.

While co-accessibility provides insight into enhancer-gene relationships, it does not directly reveal the TFs which provide regulatory input into the enhancer-promoter circuitry. We therefore performed motif enrichment analysis within snATAC-seq peaks to infer TF activity across cell types. As expected, motifs for lineage-defining TFs were enriched in a cell type-specific manner, including p53/63 and Grhl in epithelia, Gata in mesenchyme, Sox in neural crest, and Myod*/*Myog in paraxial mesoderm (Fig. 3D). However, when considered within sub-populations, we noted that p53/63 motifs were enriched in basal epithelia (cluster 6), Grhl in periderm (cluster 5), Gata in early mesenchyme (clusters 7,8,15), Nfi/Twist in late mesenchyme (clusters 9,11), Six in early paraxial mesoderm (cluster 18) and Myod/g in late paraxial mesoderm (cluster 16), consistent with the known developmental roles of these TF families (Fig.3D).

To improve TF prediction within large families that share similar binding motifs, we correlated motif accessibility with TF gene expression across joint modalities. Among other TFs, this integrative approach identified master lineage regulators including: Myod1, Myog, Myf5*/6*, Gata3/6, and P63 (Fig. 3E). Accordingly, we spatially mapped key TFs from epithelia (*Trp63*, *Grhl3*), mesenchyme (*Gata6*, *Tbx15*), paraxial mesoderm (*Six1*, *Myf5*) and neural crest (*Sox10*, *Dlx2*) (Fig. 3F,G), confirming spatial co-expression of lineage markers within their predicted cell type. Notably, *Gata6* and *Tbx15* exhibited mutually exclusive spatial domains; *Gata6* was restricted to the ventral mesenchyme and midgut (endoderm), while *Tbx15* localised to the dorsal mesenchyme (mesoderm), consistent with their distinct embryonic origins^19,20^ (Fig. 3F). Together these data establish a spatially resolved single-cell multiome atlas of embryonic skin development, which can be used as a framework to explore the complex gene-regulatory networks (GRNs) underlying skin development.

### Characterising epithelial heterogeneity at single cell resolution

As the molecular events that drive delamination of basal cells to form periderm are intrinsic to the ectoderm^5^, we subsequently focussed on characterising the epithelial compartment (Fig. 4A). Joint modality sub-clustering revealed three transcriptionally distinct epithelial populations, which were identified as mammary placode (*Nrg3*, *Pthlh*, *Edar*), basal cells (*Trp63*, *Wnt3a*, *Jag2*) and periderm (*Gabrp*, *Nectin4*, *Tacstd2*) using well characterised marker genes (Fig. 4A-C) (Supplementary Table 4).

**Fig. 4.**
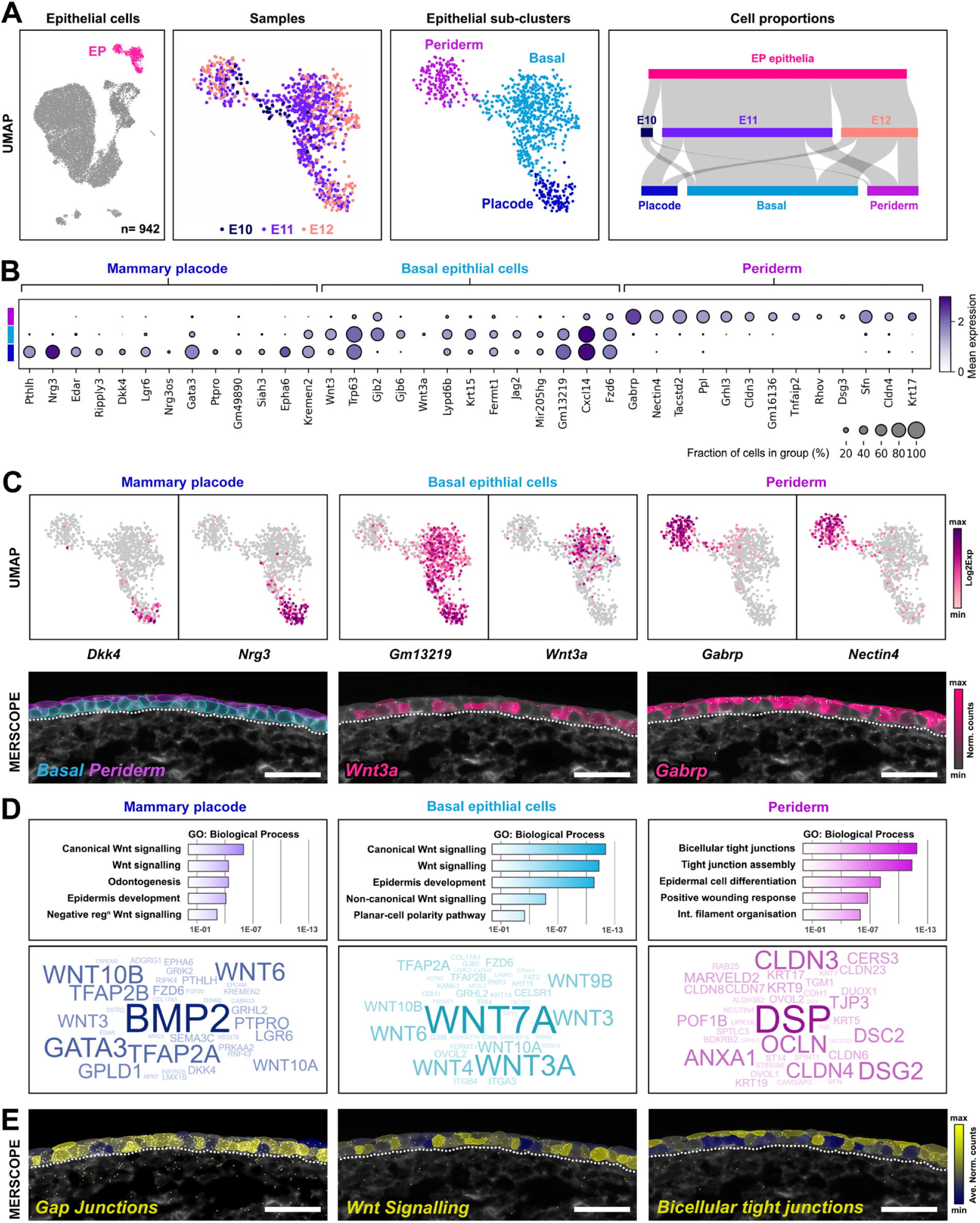
Characterisation of epithelial heterogeneity at single cell resolution. (**A**) Joint clustering UMAP of epithelial cells (EP, n= 942), followed by sub-clustering reveals three epithelial sub-populations (placode, basal cells, periderm) and their contribution from each library. (**B,C**) Marker genes and expression UMAPs of genes enriched for each sub-population. Spatial imaging (MERSCOPE) showing manual segmentation of epithelial cells (basal, periderm) and expression of *Wnt3a* and *Gabrp* (normalised transcript counts) within epithelia. (**D**) Gene ontology terms representing genes enriched in epithelial sub-clusters and corresponding gene cloud (gene frequency). (**E**) Spatial imaging of average normalised transcript counts of genes associated with ‘Gap Junctions’ (*Gjb2*, *Gjb6*) ‘Wnt signalling’ (*Wnt3a*, *Wnt7a*, *Wnt9b*) and ‘bicellular tight junctions’ (*Cldn6*, *Cldn4*, *Cldn3*, *Pof1b*, *Ocln*). Scalebars 50 µm

Spatial imaging of basal and periderm markers revealed the (largely) cell type-specific separation of transcripts. However, due to the physical proximity of these epithelial layers, unsupervised segmentation using Cellpose was limited in accurately resolving cell boundaries. To overcome this constraint, we manually segmented the surface epithelium into periderm and basal cells using multiple boundary markers and knowledge of periderm morphological features (Fig. 4C). This supervised approach was validated using snRNA-seq data, confirming that, for example, *Wnt3a* expression was restricted to basal cells, whilst *Gabrp* expression was periderm-specific (Fig. 4C).

To further define the functional identity of each epithelial sub-population, we performed Gene Ontology (GO) enrichment analysis on the top 100 marker genes per cluster. Basal cells showed enrichment in genes associated with the ‘Wnt signalling pathway’ (*Wnt6, Wnt10a, Wnt10b, Wnt3a, Wnt7b, Fzd6, Wnt9b, Wnt7a, Wnt3, Wnt4*), components of the basement membrane ‘lamina lucida’ (*Col17a1, Bcam, Itgb4, Lama3, Krt14, Lamc2, Krt5*), gap junctions (*Gjb2*, *Gjb6*) and disease associations with ‘epidermolysis bullosa’ (*Col17a1, Itgb4, Itga3, Krt14, Lamc2, Krt5, Exph5*). Similarly, mammary placode cells also showed over-representation of Wnt signalling components (*Wnt6, Wnt3, Wnt10a, Wnt10b, Fzd6, Gata3*) (Fig. 4D, Supplementary Table 5). In contrast, periderm cells were enriched in genes related to ‘bicellular tight junction assembly’ (*Cldn6, Cldn4, Ocln, Cldn3, Marveld2, Pof1b, Cldn8, Cldn23, Cldn7*), ‘cell-cell junction organisation’ (*Dsp, Ocln, Marveld2, Cdh1, Dsg2, Tjp3*), and the disease term ‘popliteal pterygium syndrome’ (*Sfn, Grhl3, Irf6*). In support of these observations, spatial mapping of gap junction (*Gjb2*, *Gjb6*) and Wnt transcripts (*Wnt3a*, *Wnt7a*, *Wnt9b*) were enriched in basal cells, whilst bicellular tight junction components (*Cldn6*, *Cldn4*, *Cldn3*, *Pof1b*, *Ocln*) were spatially enriched within periderm (Fig. 4E). Together, these data reveal a transcriptionally and spatially resolved view of epithelial differentiation in early skin development.

### P63 orchestrates periderm formation through a TF cascade

Next, we explored the transcriptional regulatory networks driving epithelial development in more detail. Focussing on epithelial cells and using a similar multimodal approach integrating gene expression and motif enrichment, we identified the activity of several key TFs as highly correlated to gene expression including: Trp63, Tead3/4, Grhl1/3; Klf4/6, Irf6 and Sox9 (Fig. 5A).

**Fig. 5.**
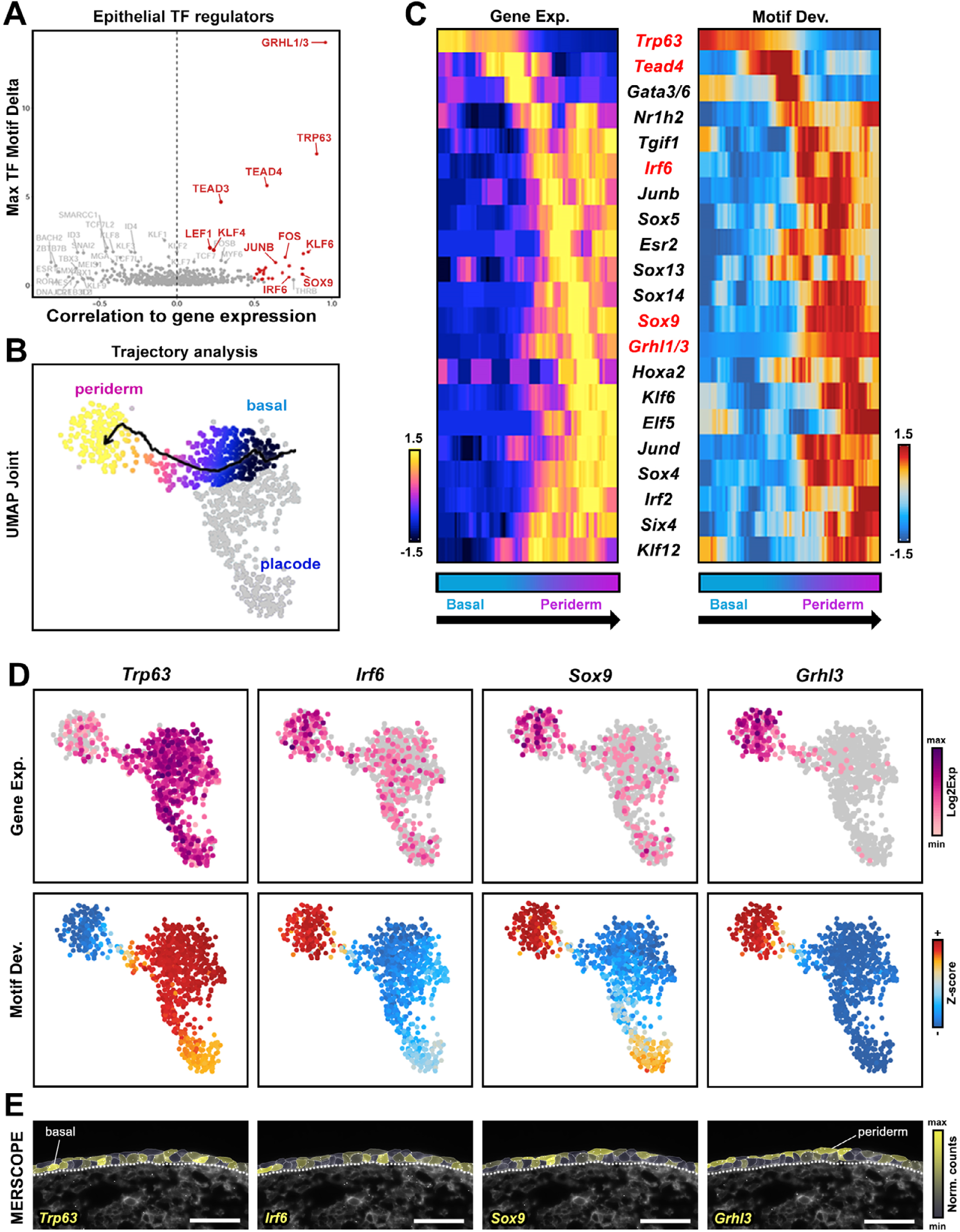
Integrative pseudotime analysis reveals transcription factor cascade underlying periderm formation. (**A**) TFregulator analysis on multimodal data identifies the transcription factors (TFs) driving epithelial development. (**B**) Pseudotemporal reconstruction of epithelial dynamics shows a trajectory from basal cells to periderm. (**C**) Integrative pseudotime analysis (gene expression correlated with motif deviation) along the trajectory uncovers a TF signalling cascade from basal cells to periderm. (**D**) Expression and motif deviation UMAPs for key TFs involved in epithelial development. (**E**) Spatial imaging (MERSCOPE) of *Trp63*, *Irf6*, *Sox9*, and *Grhl3* transcripts (normalised counts) within basal cells and periderm. Scalebars 50 µm

To uncover the TF dynamics involved in driving temporal cell state changes in epithelia, we performed multimodal integrative pseudo-time analysis along a basal cell to periderm trajectory (Fig. 5B). Through pseudo-temporal ordering, we identified a TF signalling cascade initiated by p63 in basal cells, leading to successive activity of Tead4, Gata3/6, Irf6, Sox5/9/14 and Grhl1/3 factors as cells transitioned towards periderm (Fig. 5C). Notably, down-regulation of *Trp63* correlated with decreased motif accessibility at the transition from basal cells to periderm, concomitant with increased TF activity for Tead4, Gata3, Irf6, Sox9 and Grhl3 (Fig. 5C,D). Spatial expression of these TFs supported these findings, with *Trp63* and *Irf6* transcripts concentrated within basal cells, whilst *Sox9* and *Grhl3* were enriched in periderm (Fig. 5E). Together, these data define a hierarchical transcriptional network where P63 functions as a central regulatory node, acting as a molecular gatekeeper of the cell fate transition from basal cells to periderm.

### Modelling periderm development using enhancer-gene regulatory networks (eGRNs)

To dissect the transcriptional mechanisms driving periderm formation, we constructed enhancer-gene regulatory networks (eGRNs) for key TFs implicated in the basal to periderm transition (Trp63, Irf6, Grhl). Using co-accessibility inference and gene expression correlation, we scanned epithelial-specific enhancer regions (13,874 peak-to-gene interactions, correlation >0) for the presence of TF binding motifs and linked these enhancers to putative target genes.

Construction of the P63-driven eGRN identified 4,909 predicted *cis*-regulatory interactions connecting to 2,485 target genes (Extended Data Fig. 5; Supplemental Table 6). Among the most significant targets were *Cdh1*, *Krt5*, *Krt14*, *Krt15*, *Ripk4* and *Trp63* itself. Gene set enrichment analyses confirmed the central role of P63 in regulating epithelial development, with associated target genes enriched in terms such as ‘*hemidesmosome assembly’* (*Col17a1*, *Lamb3*, *Itgb4*, *Krt14*, *Itga6*), ‘*Wnt signalling’* (*Wnt10a*, *Wnt3a*, *Jup*, *Fzd6*, *Fzd10*), ‘*epidermolysis bullosa’* (*Dsp*, *Cd151*, *Plec*, *Itga3*, *Lambc2*) and ‘*popliteal pterygium syndrome’* (*Ripk4*, *Sfn*, *Irf6*, *Grhl3*).

Downstream of P63, the predicted Irf6 eGRN (722 IRF6 *cis*-regulatory interactions to 625 target genes) (Extended Data Fig. 6; Supplemental Table 7) reflected *Irf6*’s role in promoting epithelial and periderm differentiation. Notable predicted targets included *Zfp750*, *Krt17*, *Prom2*, *Esrp2*, *Tacstd2*, *Tfap2c*, *Grhl2*, and *Irf6* itself. Gene set enrichment analyses highlighted functions in ‘tight junction assembly’ (*Cldn3*, *Cldn4*, *Cldn6*, *Cldn23*, *Patj*, *Marveld2*, *Ocln*, *Grhl2*), ‘epidermal development and differentiation’ (*Irf6*, *Krt9*, *Krt19*, *Krt15*, *Krt17*, *Spint1*, *Grhl3*), and ‘impaired skin barrier function’ (*Rarg*, *St14*, *Prss8*, *Ovol1*, *Grhl3*).

Similarly, the Grhl3 eGRN (3,971 *cis*-regulatory interactions to 2,337 target genes) (Extended Data Fig. 7; Supplemental Table 8) further supported its role in periderm specification. Consistent with known genetic interactions, Grhl3 shared multiple predicted targets with both p63 and Irf6 networks, including *Trp63*, *Irf6*, *Sfn*, *Esrp2*, *Ovol2*, *Krt5*, *Krt14*, *Krt15*, *Wnt3a*, and *Fzd6*. The Grhl3 network was enriched for pathways in ‘tight junction assembly’ (*Patj*, *Cldn1*, *Grhl2*, *Cldn6*, *Ocln*, *Dsg3*), ‘Wnt signalling’ (*Rarg*, *Jup*, *Wnt3a*, *Fzd6*, *Fzd10*, *Gata3*), and ‘impaired skin barrier function’ (*Cers3*, *Rarg*, *St14*, *Jup*, *Kdf1*, *Grhl3*).

Collectively, these interconnected GRNs reveal a tightly regulated transcriptional hierarchy in which p63, Irf6, and Grhl3 coordinate overlapping sets of target genes to orchestrate periderm formation and epithelial differentiation.

### Generation and characterisation of Trp63^-/-^ spatial multiome atlas

Recognising that eGRN predictions based solely on multiome correlations can overestimate functional enhancer-gene relationships, we sought to refine these networks using *in vivo* perturbation data. Given P63’s central role in the transcriptional cascade, we generated a complementary spatial multiome atlas from *Trp63*^-/-^ embryos at E10.5, E11.5, and E12.5. Across 12,706 cells, we profiled an average of 3,408 genes and 8,022 UMIs per cell for snRNA-seq, and detected 338,386 accessible chromatin regions. Following the same analytical pipeline applied to wild-type tissue, we first analysed *Trp63*^-/-^ data independently to infer feature linkages (Extended Data Fig. 1G-L), before integrating both datasets.

Joint modality clustering of wild-type and *Trp63*^-/-^ embryos (29,567 cells) revealed that broad cell types were preserved in *Trp63*^-/-^ skin with most co-segregating with wild-type clusters (Fig. 6A,B). However, within the epithelial compartment *Trp63*^-/-^ cells formed a distinct cluster, separate from basal, periderm, or placode populations (Fig. 6B), suggesting altered epithelial identity in the absence of P63.

**Fig. 6.**
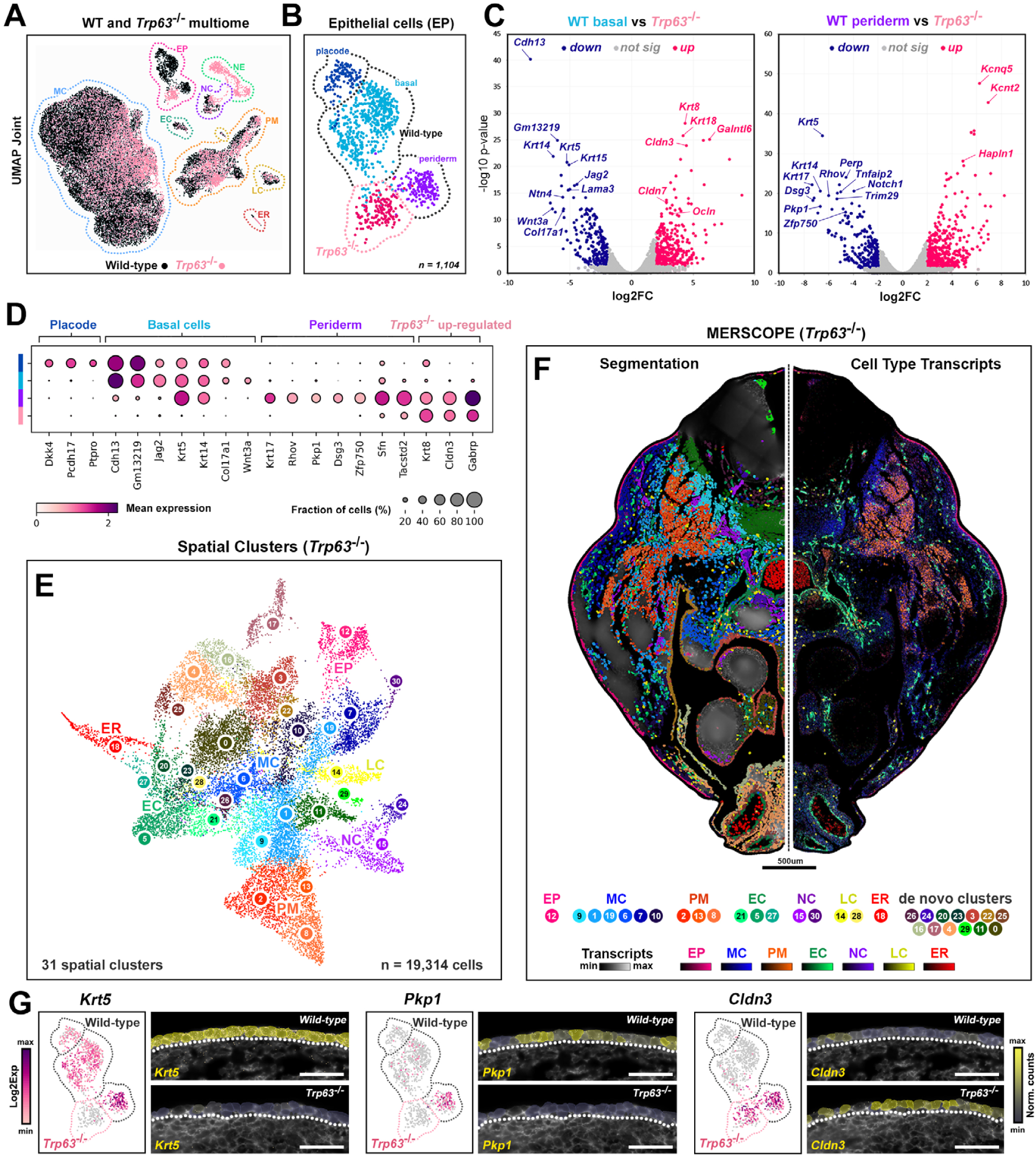
Single cell multiome and spatial analyses of *Trp63^-/-^* embryos. (**A,B**) Integration and joint clustering of wild-type and *Trp63*^-/-^ multiome (n= 29,567 cells) identifies a unique *Trp63*^-/-^ epithelial cell cluster. (**C,D**) Volcano and dotplots show the top differentially expressed genes between wild-type and *Trp63*^-/-^ epithelial subclusters. (**E-H**) Spatial imaging (MERSCOPE) of wild-type (**E,F**; sample 2) and *Trp63*^-/-^ (**G,H**; sample 4) transverse tissue sections (mirrored along dotted line) reveal spatial organisation of cell subpopulations, through segmentation, average transcript density and UMAP clustering of cell (sub)types. (**I**) Verification of gene expression changes using snRNA-seq and spatial *in situ* imaging. Scalebars 50 µm

To further characterise the *Trp63*^-/-^ epithelial state, we performed pairwise differential gene expression analyses between epithelial sub-clusters (Fig. 6C,D; Supplementary Table 9). Genes down-regulated in *Trp63*^-/-^ cells were previously associated with p63 activity, including ‘*Wnt signalling’* (*Wnt10b*, *Lgr5*, *Dkk4*, *Wnt5a*, *Kremen2, Wnt3a, Wnt9b, Fzd6, Fzd10*), ‘*placode specification’* (*Nrg3*, *Ripply3*, *Pthlh*, *Edar*, *Edaradd)*, ‘*epidermis development’* (*Krt5*, *Krt14*, *Krt15*, *Lama3*, *Col17a1*), ‘*cell junctions’* (*Kirrel3*, *Notch1*, *Jag1*, *Pof1b*, *Trim29*, *Perp*) ‘*cornified envelope’* (*Dsg2*, *Dsg3*, *Pkp1*, *Jup*, *Tgm1*) and the disease term ‘*popliteal pterygium syndrome’* (*Sfn*, *Grhl3*, *Irf6*), correlating well with predicted targets identified from the P63 eGRN. Conversely, genes characteristic of simple epithelia, such as *Krt8* and *Krt18*, were up-regulated in *Trp63*^-/-^ cells. However, these were co-expressed with markers of both basal (*Wnt3*) and periderm lineages (*Cldn3*, *Cldn7*, *Ocln*, *Gabrp*, *Tacstd2*), suggestive of an aberrant cell state and a loss of cell identity (Fig. 6D).

To verify gene expression changes *in situ*, we generated three additional spatial transcriptomics datasets from E12 *Trp63*^-/-^ tissue sections (Extended Data Fig. 8 and 9; samples 4-6). Using our established spatial analysis pipeline, we detected a further ∼3 million transcripts within 52,390 segmented cells. Comparing spatial gene expression in wild-type (sample 2) and *Trp63*^-/-^ tissue (sample 4) (Fig. 6E-G), we confirmed the altered expression of key epithelial genes identified by snRNA-seq (Fig. 6G).

### Integrated genome-wide analysis of P63 enhancer-gene targets using Trp63^-/-^ mice

Next, we assessed how loss of p63 affected the accessible chromatin landscape. While promoter accessibility remained largely unchanged between wild-type and *Trp63*^-/-^ embryos, we readily identified distal epithelial enhancers with reduced accessibility, many demonstrating P63 occupancy in epithelial tissue (P63 ChIP-seq, E11.5 face) and/or contained predicted P63 motifs. Furthermore, loss of chromatin accessibility at these enhancers often coincided with a loss of predicted enhancer-gene interactions relative to wild-type data, supporting their functional significance.

To refine p63 enhancer-gene targets genome-wide, we integrated P63 ChIP-seq (E11.5 face, 10,209 peaks) with *Trp63*^-/-^ multiome data to identify P63-bound loci associated with reduced enhancer accessibility, down-regulated gene expression, and loss of enhancer-gene interactions when compared to the wild-type p63 eGRN. This integrative approach yielded a refined network comprising 1,471 putative p63 *cis*-regulatory interactions targeting 910 genes. To prioritise the most functionally relevant targets, we intersected this dataset with significantly down-regulated epithelial genes identified from *Trp63*^-/-^ pairwise expression analyses (fc<-1, p<0.05), resulting in a high-confidence subset of 626 interactions targeting 354 genes (Supplementary Table 10).

As verification of our strategy, we profiled the snATAC-seq signals of these regions in wild-type and *Trp63*^-/-^ epithelial cells together with P63 occupancy, confirming significantly reduced accessibility in regions where P63 can bind (Fig. 7A). Subsequently, we visualised this high confidence p63 regulatory network as a directed graph (Fig. 7B), revealing established basal epithelial (e.g. *Krt5*, *Perp*, *Krt14*, *Krt15*, *Trim29*), Wnt (e.g. *Wnt3a*, *Wnt9b*, *Fzd6*), and novel targets such as the atypical cadherin *Cdh13 (T-cadherin)* and long non-coding RNA (lncRNA) *2610035D17Rik* (*Rason; LINC00673*). Closer examination of the *Cdh13* locus revealed five intragenic p63-dependent putative enhancers spanning nearly 900 kb (Fig. 7C), that exhibited reduced accessibility, disrupted chromatin interactions and significantly down-regulated gene expression in *Trp63*^-/-^ epithelia (Fig. 7D). Similarly, we identified three p63-dependent enhancers (- 58 kb, -30 kb, +69 kb) linked to the novel target *2610035D17Rik*, which correlated with down-regulated expression in *Trp63*-/- epidermis (Fig. 7E,F).

**Fig. 7.**
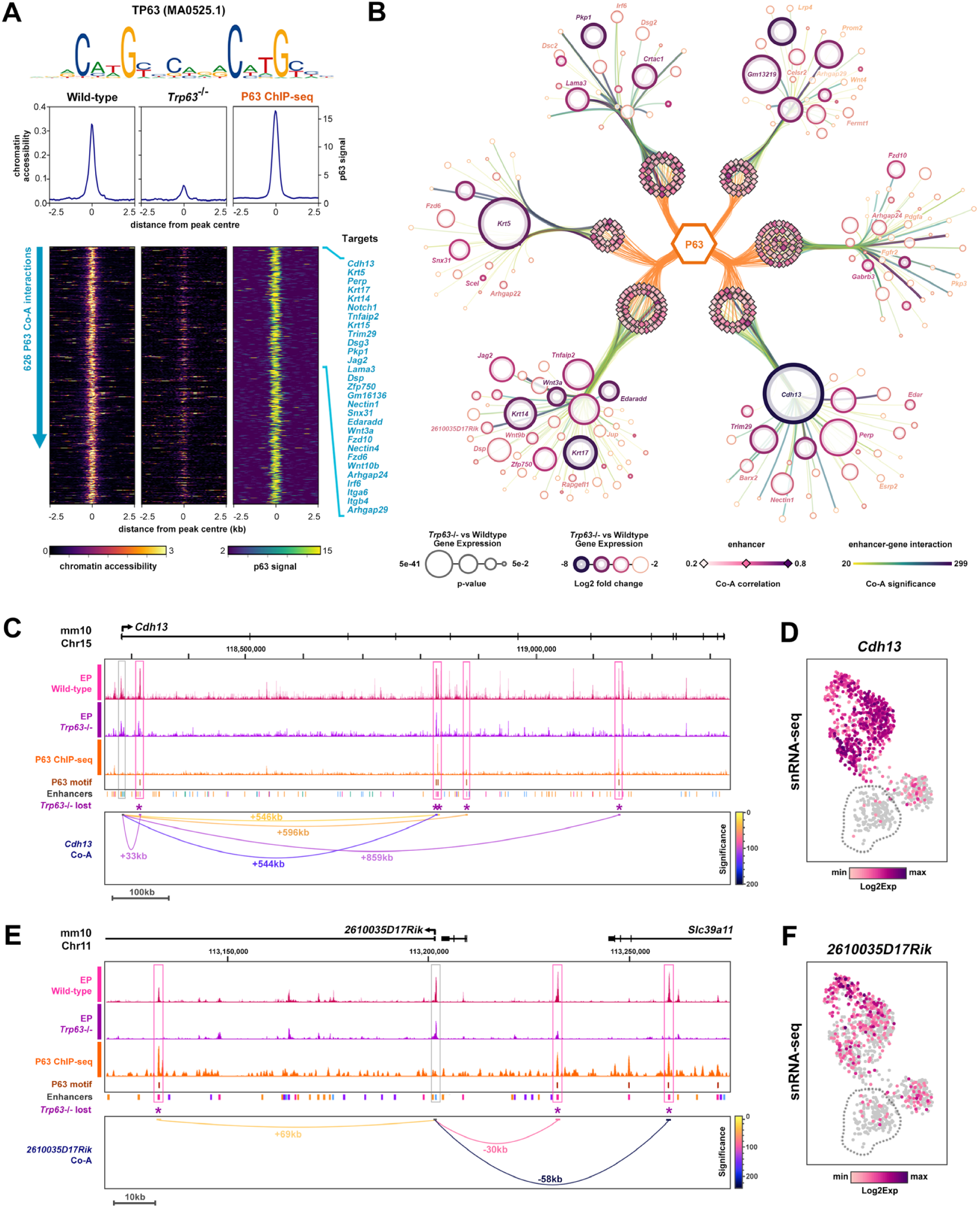
Integrated multiomics deciphers the P63-dependent enhancer-gene regulatory networks. (**A**) snATAC-seq signals of 626 P63-dependent enhancers in wild-type and *Trp63*^-/-^ epithelia, correlated with P63 occupancy in E11.5 facial epithelia (ChIP-seq). (**B**) P63 enhancer-gene regulatory network graph of high confidence targets that are significantly downregulated in *Trp63*^-/-^ epithelia. (**C-E**) Long range P63-dependent enhancers linked to *Cdh13* (**C**) and *2610035D17Rik* (**E**), demonstrate reduced chromatin accessibility and loss of linkage to gene promoter. Expression of *Cdh13* (**D**) and *2610035D17Rik* (**F**) (snRNA-seq) in wild-type and *Trp63*^-/-^ epithelia.

### P63 orchestrates epithelial cell fate by regulating key signalling pathways and downstream transcription factors

Integrated gene expression analyses indicated Wnt and Eda signalling pathways were significantly disrupted in the epithelia of *Trp63*^-/-^ mice, supporting gene regulatory network predictions and corroborating earlier reports^21^. Accordingly, several key components of these pathways, including Wnt ligands (*Wnt3a*, *Wnt9b*, *Wnt10b*), Frizzled receptors (*Fzd6*, *Fzd10*), the non-canonical Wnt receptor (*Celsr2*), and Eda pathway components (Edar, Edaradd), were identified as candidate direct targets (Fig. 7B). For each target, we confirmed differential enhancer accessibility and validated altered epithelial expression by snRNA-seq and *in situ* spatial transcriptomics (Extended Data Fig. 10). For example, three P63-bound enhancers associated with *Fzd6* (promoter, -28 kb, -32 kb) exhibited reduced chromatin accessibility, loss of regulatory linkage, and decreased transcript levels in *Trp63*^-/-^ epidermis (Extended Data Fig. 10A-C).Similarly, P63-dependent enhancers were linked to down-regulated expression of *Fzd10* (+49 kb), *Wnt3a* (+6 kb), *Edaradd* (+34 kb), *Edar* (+34 kb) and *Celsr2* (+13 kb, -9 kb, -66 kb) in *Trp63*^-/-^ epidermis (Extended Data Fig. 10).

P63 also occupied enhancers linked to several key TFs with roles in epithelial remodelling, differentiation and barrier formation^22–27^, including *Ovol2* (+50 kb, +56 kb), *Zfp750* (-3.3 kb, -39 kb), *Mafb* (+57 kb, +111 kb, +218 kb), *Barx2* (-98 kb, +183 kb), *Tead4* (+ 38kb), *Mycl* (-50 kb), *Irf6* (-12 kb) and *Grhl3* (+0.4 kb, +3 kb), Extended Data Fig. 11; Extended Data Fig. 12A-C). Apart from Tead4, reduced accessibility/linkage of P63-bound enhancers correlated with down-regulated expression of their predicted target gene in *Trp63*^-/-^ epidermis. Together with pseudo-temporal modelling placing the activity of several of these targets downstream of P63, our data support a role for P63 in establishing a complex TF cascade to coordinate changes in epithelial cell fate, periderm formation and epithelial differentiation.

### P63-dependent enhancers regulate critical genes implicated in peridermopathies

Failure of periderm formation and/or differentiation results in a spectrum of congenital conditions collectively termed ‘peridermopathies’^7–12,28^. Studies in humans, mice, and zebrafish have revealed a complex transcriptional network essential for periderm development, involving genes such as *Irf6*, *Grhl3*, *Prkci*, *Ripk4*, *Jag2*, *Chuk*, *Sfn*, and *Kdf1*. Notably, *Trp63*^-/-^ embryos fail to form normal periderm and we identified P63-dependent enhancers associated with all these genes, except *Chuk* (*Ikka*), highlighting the pivotal regulatory role of P63 within this network.

Complementing our findings with *Irf6* (-12 kb) and *Grhl3* (+0.4 kb, +3 kb), our data also revealed P63-bound elements co-accessible with *Ripk4* (+4 kb, +12 kb), *Prkci* (+4 kb), *Jag2* (+1 kb, +44 kb, +48 kb), *Sfn* (-3 kb), and *Kdf1* (+86 kb) (Extended Data Fig. 12). Apart from *Prkci* and *Kdf1*, reduced enhancer accessibility in *Trp63*^-/-^ epidermis correlated with down-regulated gene expression. Interestingly, one P63-dependent enhancer was co-accessible with both *Sfn* (-3 kb) and *Kdf1* (+86 kb) promoters (Extended Data Fig. 12G), epigenetically linking two genes that not only share phenotypes in knockout mouse models but have also been shown to interact genetically during epidermal development^29^.

To validate the novel P63-dependent putative enhancers identified from our multiomic analyses, we employed CRISPR interference (CRISPRi) in human keratinocytes. For comparative analyses, we lifted over conserved *cis*-regulatory enhancer-gene interactions to the human genome (UCSC: hg19), retaining 75% of epithelial interactions, and integrated with P63 occupancy data^30^ and histone methylation and acetylation profiles from primary keratinocytes^31^ to support enhancer activity (Fig. 8; Extended Data Fig. 13). Using HaCaT cells stably expressing *dCas9-KRAB-MeCP2* (dCas9-KM2)^32^, we targeted individual regulatory elements with up to four sgRNAs per enhancer. Target gene expression was quantified by qPCR relative to a scrambled sgRNA control (SCR2), with *RAB1A* serving as a positive control^33^.

**Fig. 8.**
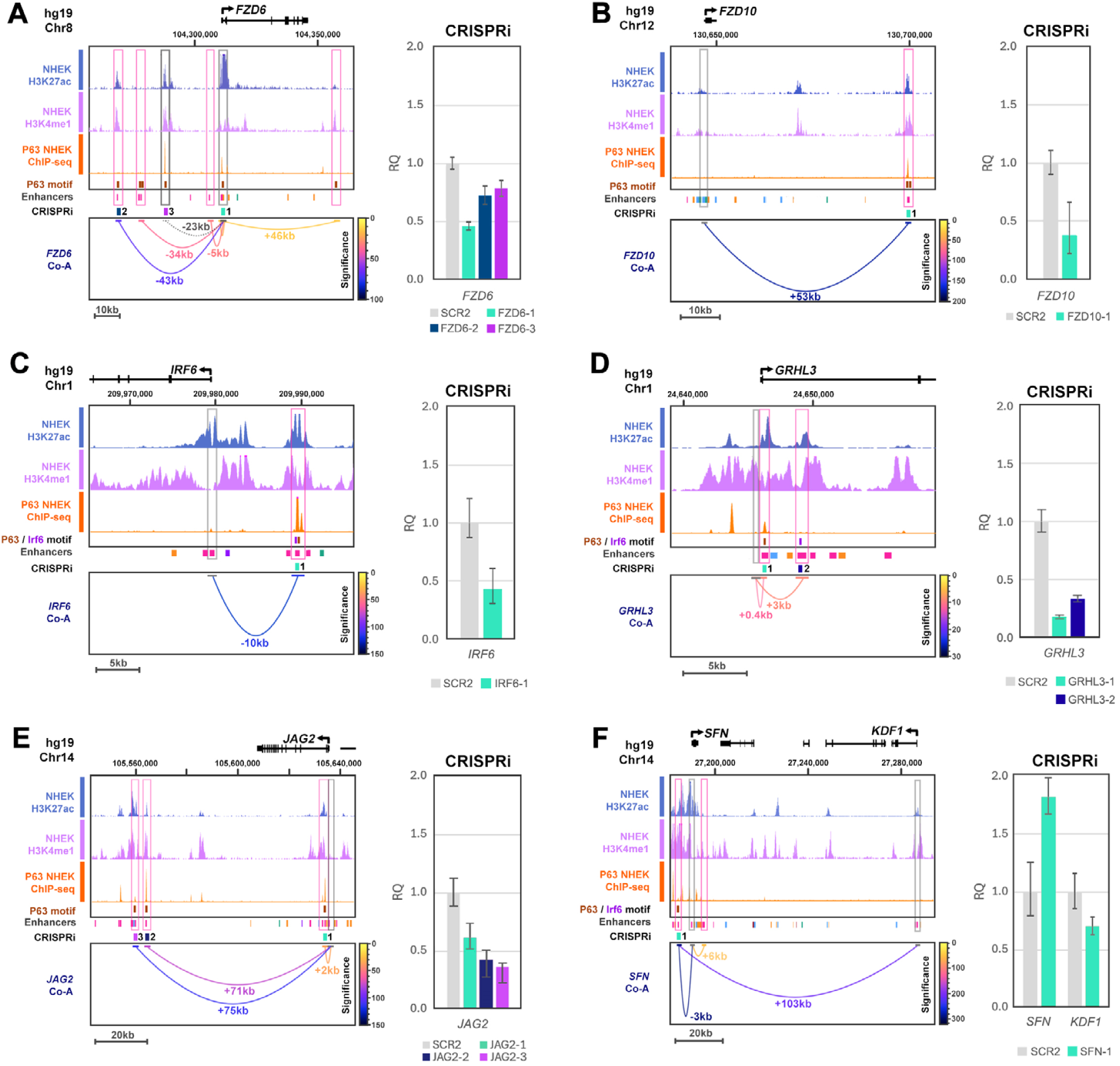
Comparative validation of P63 enhancer-gene targets. **(A-F)** P63-dependent enhancer-gene targets in human (hg19), targeted by CRISPRi in HaCaT cells. UCSC views of comparative genomic regions for *FZD6* (**A**), *FZD10* (**B**), *IRF6* (**C**), *GRHL3* (**D**), *JAG2* (**E**) and *SFN* / *KDF1* (**F**), together with ENCODE histone methylation (H3k4me1, H3k27ac) and P63 occupancy (ChIP-seq^30^), show enhancers targeted by CRISPRi followed by qPCR profiling of target gene expression in cells expressing enhancer guideRNAs versus scrambled controls (SCR2). Error bars denote RQ expression min/max confidence intervals (95%).

Focusing on the Wnt signalling pathway, we identified conserved P63-bound enhancers marked by active histone modifications and associated with both canonical (*WNT3A*, +6 kb; *FZD6*, promoter, -43 kb, -23 kb; *FZD10*, +53 kb) and non-canonical (*CELSR2*, +9 kb, -14 kb) targets (Fig. 8A,B; Extended Data Fig. 13). Targeted repression of enhancers linked to Wnt machinery (*FZD6*, *FZD10, WNT3A*) led to a reduction in transcript levels (*FZD6* -1.3 to -2.2 fold; *FZD10* -2.6 fold; *WNT3A* -1.7 fold). Moreover, repression of elements associated with *CELSR2* resulted in robust reduction in transcript levels (-2.9 to -3.6 fold). Together, these findings support a functional role for P63-bound regulatory elements in modulating components of the Wnt signalling machinery.

Next, we explored enhancers associated with the periderm GRN by targeting conserved elements linked to *IRF6* (-10 kb), *GRHL3* (+0.4 kb, +3 kb) and *PRKCI* (+4 kb) using CRISPRi (Fig. 8D-F; Extended Data Fig. 13). Repression of the P63 occupied *IRF6*-linked element (-10 kb) reduced *IRF6* transcripts by -2.3 fold, validating our enhancer-gene predictions and reinforcing previous studies^15–17^. Similarly, repression of *GRHL3*-linked enhancers containing P63 (+0.4 kb) or IRF6 binding sites (+3 kb) led to a robust reduction in transcript levels (-1.3 to -1.9 fold) reinforcing the regulatory interplay between these TFs^34^ (Fig. 8C,D). In contrast, targeted repression of the *PRKCI*-linked element (+4 kb) did not significantly alter transcript levels (Extended Data Fig. 13).

We also functionally assessed conserved enhancers associated with epidermal/periderm differentiation targets *JAG2* (+2 kb, +71 kb, +75 kb) and *SFN* / *KDF1* (-3 kb / +103 kb) (Fig. 8E,F). Analysis of the *JAG2* locus revealed a cluster of P63-bound enhancers, marked by active histone modifications and at considerable distance from the promoter. Targeted repression of individual *JAG2*-linked enhancers resulted in reduced transcript levels (-1.6 to -2.7 fold) confirming P63 positively regulates Jag2-Notch1 signalling^35–37^ (Fig. 8E). As *SFN* and *KDF1* are syntenic and co-accessible with the same P63-bound enhancer, following CRISPRi we quantified transcripts of both genes. Remarkably, repression of this enhancer increased *SFN* transcript levels (+1.8 fold increase) while reducing *KDF1* transcripts (-1.4 fold), suggesting the element directly regulates *KDF1*, with secondary effects on *SFN* (Fig. 8F). Together with reports that *Kdf1* and *Sfn* genetically interact to regulate epidermal proliferation/differentiation^29^, these findings point to a more complex regulatory relationship that warrants further study.

## Discussion

The transition from a simple single-layered ectoderm to a stratified, multi-layered epidermis is a key milestone in vertebrate skin development, which begins when basal epithelial cells detach from the basement membrane to produce a transient superficial layer known as the periderm. While the transcriptional and chromatin dynamics of embryonic skin development have been studied at early (E9) and later (E12.5 - E18.5) stages^21,38–40^, the regulatory mechanisms surrounding periderm formation are poorly understood. Here, we detail the first comprehensive, spatially resolved single cell multiome atlas of early embryonic skin development (E10 - E12), capturing the transcriptional and epigenetic programs underpinning epidermal lineage diversification and periderm formation.

Through joint modality profiling of over 21,000 nuclei from lateral skin and sub-cellular *in situ* spatial imaging, we spatially resolved the major embryonic lineages, including epithelia, mesenchyme, paraxial mesoderm, endothelia and neural crest derivatives, and further characterised 18 transcriptionally distinct sub-populations. Spatial transcriptomics enabled these cell populations to be mapped at high resolution *in situ*, revealing their precise anatomical organisation, and uncovered *de novo* populations beyond the scope of tissue microdissection.

By leveraging joint gene expression and chromatin accessibility measurements within the same cell, we were able to infer TF activity with greater confidence than expression or motif enrichment alone. This integrative approach allowed us to identify functionally relevant TFs underlying lineage commitment, for example *Myod1*, *Myog* and *Myf5*/6, driving skeletal muscle and myogenesis^41^. These findings illustrate the power of multimodal analyses to infer gene regulatory programs driving cell fate specification.

Focussing on the epithelial compartment, our spatial single-cell analyses revealed distinct transcriptional signatures corresponding to basal and periderm sub-populations. Spatial mapping verified the basal localisation of canonical Wnt pathway components, while genes associated with tight junctions and differentiation were enriched in periderm. These findings reinforce the role of Wnt/Edar signalling in appendage specification^21,42,43^ and the importance of periderm-specific genes in promoting epithelial polarity, differentiation and barrier formation^2,39^.

Our data provide compelling evidence that P63 is a central node in orchestrating periderm formation and epithelial differentiation. Reconstruction of the TF dynamics underlying the basal to periderm transition revealed a hierarchical TF cascade (Tead4, Irf6, Grhl3, Sox9) initiated by P63 in basal cells. This cascade aligns with previous studies, implicating several key TFs at the interface between epithelial proliferation/differentiation fate decisions and are essential to periderm integrity^6,22,23,25,34,44,45^. Intriguingly, our data also reveal the novel transcriptional activity of several members of the Sox family associated with periderm, particularly *Sox9*, suggesting a potential role in periderm differentiation. Further studies will be required to elucidate the functional contribution of the Sox family to periderm development.

Using joint modality correlations within the same cell, we identified over 120,000 significant peak-to-gene interactions across all cell types, providing a framework for exploring gene regulatory mechanisms underlying cell fate specification of several lineages. Within epithelia, we reconstructed enhancer-gene regulatory networks for key TFs in the cascade (P63, Irf6, Grhl3), revealing a complex interconnected network of co-regulated genes that converge on Wnt signalling, epithelial barrier formation and pathways implicated in congenital epithelial disorders. Integration with *Trp63*^-/-^ multiome data further supported these predictions, showing reduced chromatin accessibility at distal enhancers, disruption of enhancer-gene interactions, and downregulation of key genes required for epithelial differentiation and periderm formation. Importantly, our dataset also provides a rich resource for exploring novel P63 enhancer-gene targets, including candidates such as *Cdh13* and *2610035D17Rik*, whose function in epidermal development remain unexplored.

Our enhancer-gene regulatory modelling also identified P63-dependent enhancers associated with key peridermopathy genes, including *Grhl3*, *Irf6*, *Ripk4*, *Sfn*, *Jag2*, and *Kdf1*, expanding the transcriptional influence of P63 within this network. Many of these enhancers were conserved in human, and comparative CRISPR interference validated several enhancer-gene interactions in their native genomic context. Beyond defining mechanisms of periderm formation, this regulatory atlas also provides a central resource for exploring disease-associated loci, enabling the functional interpretation of regulatory variants implicated in epithelial disorders.

In summary, our spatial single cell multiome atlas provides the most comprehensive regulatory blueprint of early skin development to date and provides a mechanistic framework for dissecting the regulatory architecture of transcriptional networks driving epithelial differentiation.

## Methods

### Mouse tissue

Generation and genotyping of *Trp63*^−/−^ mice have been described previously^46^. Mice were housed in accredited animal facilities at the University of Manchester. All procedures were approved by the University of Manchester Ethical Review Committee and are licensed under the Animal (Scientific Procedures) Act 1986. *Trp63*^−/−^ embryos were obtained by intercrossing *Trp63*^+/-^ mice, the morning of a vaginal plug being considered as E0.5.

### Histology and immunohistochemistry

Embryos for histological and immunohistochemical analyses were harvested at E10-E12.5, fixed overnight in fresh 4% paraformaldehyde, processed to paraffin wax using standard protocols and sectioned at 6 µm thickness. Sections for immunofluorescence analysis were dewaxed in xylene, rehydrated through a graded series of ethanols to dH_2_0, followed by standard antigen retrieval in 10 mm citrate buffer (pH 6.0). Primary mouse antibodies were used with M.O.M kit (Vector Laboratories, UK), while primary rabbit antibodies were diluted in 10% serum (of the same species secondary antibody is raised in), prior to detection using directly conjugated fluorescently labelled secondary antibodies (Alexa Fluor+ 488, 594; 1:500; ThermoFisher, UK). Prepared slides were mounted with Vectashield Vibrance antifade mounting media containing DAPI (Vector Laboratories, UK). Monochrome images for each channel were acquired on a Leica DMRB microscope using Tucsen FL-20BW camera, pseudo-coloured and merged using Mosaic 3.0 software (Tucsen), before importing into Adobe Photoshop CS6 to assemble figures.

### Single cell multiome sample processing, library preparation and sequencing

For single cell multiome analysis, at least three litters of embryos for each embryonic age and genotype were obtained from timed matings, harvested and dissected into ice-cold DPBS. Dorsal torso skin from the region indicated (Fig. 1A) was micro-dissected from up to ten embryos, tissue pooled and immediately processed for single cell multiome analysis using a modified 10X Genomics protocol. Briefly, the tissue was collected, resuspended and dissociated to single cells in TrypLE (Invitrogen) at room temperature, followed by gentle trituration with barrier tips to generate a single cell suspension. TrypLE was inactivated by addition of chilled DPBS and single cells isolated by filtering through FlowMI filters. Nuclei were extracted using 0.1X lysis buffer, washed and resuspended in a minimal volume of 1X nuclei buffer. Single cells and nuclei were quantified using Countess 3 cell counter (ThermoFisher, UK), before proceeding with multiome snRNA-seq and snATAC-seq library preparation and sequencing, carried out by the University of Manchester Genomic Technology Core Facility. Sequencing was performed on the Illumina NextSeq 500 platform.

### Single cell multiome data analyses

Each multiome sample was processed individually using Cell Ranger ARC v2.0.0, followed by downstream analysis and sample aggregation with ArchR v1.0.2. Genome and gene annotations were based on the mm10 reference for both Cell Ranger ARC and ArchR. As part of initial QC, doublets were identified and removed separately for the GEX and ATAC modalities. Cells with total read counts or numbers of detected genes below three absolute deviations across the three samples were also excluded. ArchR partitions the genome into 500 bp bins and applies iterative Latent Semantic Indexing (LSI) for dimensionality reduction on RNA and ATAC modalities independently, as well as jointly. These LSIs were then used for UMAP generation. Clustering was performed on the joint LSI using ArchR’s *addClusters()* function, which applies Seurat’s Louvain algorithm, and clusters were annotated using RNA modality marker genes. Peak calling was performed with ArchR’s *addReproduciblePeakSet()* function, which uses MACS2 to call peaks on pseudo-bulk replicates generated by merging similar cell types, followed by iterative removal of overlapping peaks to retain the most significant. To identify transcription factor (TF) binding sites enriched in each cluster, curated motif annotations (cisBP by default) were added, followed by calculation of motif deviations to assess accessibility relative to background. Enriched motifs were identified based on motif deviations together with cluster-specific marker peaks. For pseudotime analysis, we applied ArchR’s trajectory analysis, which uses a user-defined backbone to construct a cellular trajectory and assign pseudotime values to cells. Integrative pseudotime analysis of positive TF regulators was carried out using *correlateTrajectories()*, which identifies TFs whose motif deviations along pseudotime correlate with their gene expression. This information was visualised jointly using *plotTrajectoryHeatmap()*. Gene expression visualisation was performed using Loupe v6 and Cellxgene VIP v1.1.1.

### MERSCOPE panel design, sample preparation, processing and analyses

We designed a 285-gene panel for MERFISH (Supplementary Table 2), comprising 167 genes from snRNA-seq differential expression of cell types, and 118 genes representing TF targets, signalling pathways and genes relevant to epithelial development. We used the Vizgen Gene Panel Design Portal, ensuring >30 probes per target and assessing FPKM and sum UMIs for target genes to ensure transcript abundance did not affect sensitivity.

Tissue samples were fixed and processed to paraffin wax as previously described, sectioned at 5 µm thickness and transferred using a water bath to capture discs. Sections were dried on a slide rack for 1 hr, air dried for 30 mins and transferred to -20 °C prior to processing. The samples were processed according to Vizgen’s FFPE workflow with extended clearing steps. Briefly, samples underwent deparaffinization and decrosslinking, cell boundary staining, RNA anchoring and gel embedding, digestion (2 hr at 37 °C), clearing and photobleaching (4 hr clearing (1:4) at 47 °C followed by 20hr (1:10) at 47 °C; 3h photobleaching; 5 hr clearing (1:4) at 37 °C followed by 20 hr (1:10) at 37 °C, quenching, probe hybridisation and washing.

All sections were imaged and stitched together at 60x and transcripts decoded using the panel codebook. Raw data was initially processed on MERSCOPE Instrument software. Cell segmentation was performed using the built-in CellPose 2.0 algorithm, with transcripts assigned to cell boundaries as part of the Vizgen workflow.

Metadata and gene courts were extracted from MERSCOPE outputs to create an .AnnData object in Python. Quality control was performed, such that cells with fewer than five counts and genes expressed in fewer than ten cells were removed. Data was then normalised with the Scanpy v1.9.8 *normalize_total()* function, followed by log1p transformation and scaling to zero mean and unit variance. PCA was performed on the scaled data, and the top 20 components were used to construct a nearest-neighbour graph with a neighbourhood size of 20. This graph was used to compute UMAP projections and Leiden clustering. Spatial data was integrated with the GEX component of sc-multiome data using Scanpy’s *AnnData.concatenate()* function, with sample specified as the batch key and the intersection of genes retained. Pearson correlations between the sc-multiome and spatial datasets were then calculated using Scanpy’s *pl.correlation_matrix()* function, based on the top 50 principal components of the merged data. These correlations were used to help annotate spatial clusters.

### Enhancer-gene regulatory network (eGRN) construction

Epithelial enhancers that were co-accessible and positively correlated with gene promoters (Cell Ranger ARC) were scanned for motifs (p63, Irf6, Grhl) using MEME-suite tools using default parameters. To refine the P63 eGRN, wild-type interactions were intersected with *Trp63*^-/-^ multiome interactions, to identify enhancer-gene interactions with loss of chromatin accessibility. P63 ChIP-seq data (E11.5 face; unpublished study) was also intersected with wild-type enhancers as further evidence of P63 occupancy. DeepTools (Galaxy server) was used to profile P63 eGRN enhancers using epithelial snATAC-seq and ChIP-seq signal tracks. eGRNs were visualised as directed network graphs using Cytoscape v3.9, with interactions filtered for significance for clarity. Gene ontology analyses of eGRNs were performed using Enrichr (https://maayanlab.cloud/Enrichr).

### Comparative UCSC genome analyses

Enhancer and promoter snATAC-seq peaks were ‘lifted’ from mouse (mm10) to human (hg19) genome assemblies using UCSC liftOver tool using default parameters. Only interactions where both peak-gene enhancers were present and on the same chromosome were considered valid for exploring in human data.

### CRISPR Interference

To functionally interrogate enhancer-gene interactions, we used a HaCaT cell line stably expressing the CRISPR repressor dCas9-KRAB-MeCP2 (dCas9-KM2)^32^. Up to 4 sgRNAs were designed across each regulatory element using the web tool CRISPOR (https://crispor.gi.ucsc.edu/) and cloned into a lentiviral delivery vector co-expressing a Neomycin resistance gene (pLKO5.sgRNA.EFS.NeoR), allowing antibiotic co-selection of cells stably expressing the CRISPRi machinery and sgRNA. For each locus, the effect of on-target sgRNA on gene expression was compared to control cells targeted with scrambled sgRNA (Scramble2; SCR2)^47^ using RT-qPCR (Supplementary Table 11).

### Guide RNA design and cloning

Guide (g)RNAs (Supplementary Table 11) were designed using the CRISPOR web design tool. Complementary gRNA oligo pairs with 5′ CACC (fwd) and 5′CAAA (rev) overhangs were phosphorylated and annealed in a thermocycler at 37 °C, 30 min; 95 °C 5 min, then ramped down to 25 °C at 5 °C/min using T4 Polynucleotide kinase (NEB) and 10× T4 ligation buffer (NEB).

Transfer plasmid pLKO5.sgRNA.EFS.Neo (Genome Editing Unit, University of Manchester) was digested with FastDigest Esp3I, Fast AP, 10× Fast Digest Buffer at 37 °C, 30 min (Fermentas) followed by the ligation of the annealed gRNA duplex overnight using T4 DNA ligase (NEB) and 10× T4 DNA ligation buffer (NEB). Stbl3™ Chemically Competent E. coli (Thermo Fisher, C7373-03) were transformed with gRNA-containing pLKO5 plasmids by heat shocking for 45 s at 42 °C. Plasmid DNA from selected colonies was isolated using a SmartPure Plasmid miniprep kit (Eurogentic) and Sanger sequenced using the hU6 sequencing primer to confirm the insertion and sequences of cloned gRNAs.

For lentivirus production where more than one guide was used, equimolar amounts of each plasmid was pooled and used for packaging.

### Lentivirus production

To prepare gRNA-containing lentiviral particles, Lenti-X HEK293T cells were transfected with transfer plasmid (pLKO5.sgRNA.EFS.Neo), pMD2.G (VSV-G envelope plasmid Addgene # 12259), pMDLg/pRRE (packaging plasmid Addgene #12251) and pRSV-REV (packaging plasmid Addgene # 12253) (total 12 µg plasmid DNA diluted in 2mL Optimem, mass ratio 4:1:1:2, respectively). 36 µl of 1mg/ml stock PEI was added to each tube (volume based on a 6:1 ratio of PEI:DNA), vortexed and incubated at room temperature for 12-15 minutes, then the complexes added to the cells and incubated for 72-hours at 37°C/5% CO_2_.

Supernatant containing the viral particles was collected and cleared by centrifugation then filtered through a 0.45µm syringe filter. The lentivirus was concentrated using Lenti-X Concentrator according to the manufacturer’s protocol (Takara Biosciences), resuspended in PBS, and stored at -80 °C.

### CRISPR interference (CRISPRi) assay

For each pool of sgRNA and an untransduced control well, HaCaT-dCas9-KM2 cells were plated in 6-well TC-treated plates (Corning) at 3×10^5^ cells/well in 3 ml complete HaCaT media and cultured overnight. After 24-hours, cells were washed in Dulbecco’s phosphate buffered saline (DPBS) (Sigma) and media exchanged for 2 ml fresh media containing 24 µl 1 mg/ml Polybrene (Sigma H9268) and 20 µl concentrated lentiviral particles carrying pLKO5.sgRNA.NeoR expressing sgRNA. After 24 hrs, cells were washed and the media exchanged for 3mL complete HaCaT media. Cells were cultured for a further 24-48 hrs until wells were confluent, then transferred to T75 flasks in complete HaCaT media and incubated overnight. Media was then exchanged for complete HaCaT media + 800µg/mL Geneticin (Gibco). Selection was carried out for ∼7-10 days, with media changed every 2-3 days, until all untransduced control cells had died. Cells were then moved to a maintenance concentration of Geneticin (400µg/mL) and rested overnight before being processed for RNA.

RNA was extracted using an Qiagen RNeasy Mini Kit following the manufacturer’s protocol. To quantify changes in transcripts, we designed qPCR primers for the genes of interest and ensured melt curves produced high quality amplification of the desired product. qPCR reactions were set-up in 96-well plate format, using Power SYBR™ Green RNA-to-CT™ 1-Step Kit (Thermo Fisher Scientific, UK), according to manufacturer’s instructions. For each locus, the effect of on-target sgRNA on gene expression was compared to control cells targeted with scrambled sgRNA (Scramble2; SCR2), using the delta-delta-CT method after normalising to a housekeeping gene (*YWHAZ*). Reactions were performed in (at least) triplicate for each condition/gene.

## Acknowledgements

We should like to thank the Genome Editing Unit and Genomic Technologies Core Facilities at the University of Manchester for providing technical support and expertise. This research was supported by the BBSRC (BB/V011626/1 to MJD).

---

**Extended Data Fig. 1.**
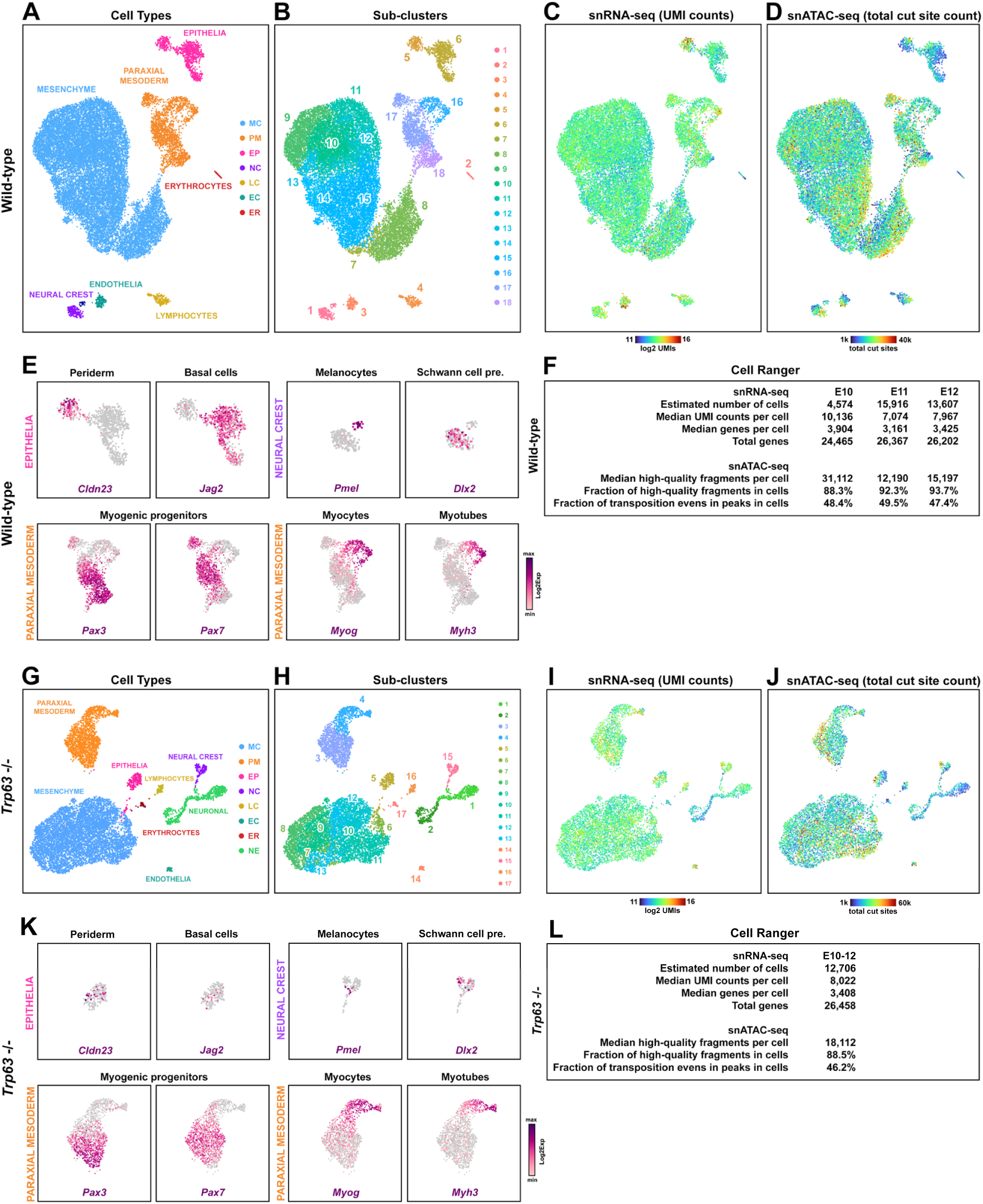
sc-multiome subclusters and mapping statistics. (**A-L**) Joint modality sc-multiome data for (**A-F**) wild-type and (**G-L**) *Trp63*^-/-^ samples, showing cell type (**A,G**), subclusters (**B,H**), UMI counts (**C,I**) and total cut site counts (**D,J**) as UMAPs. (**E,K**) Expression UMAPs for markers of epithelial, neural crest and paraxial mesoderm subclusters. (**F,L**) Cell ranger sequencing and mapping metrics for each sample.

**Extended Data Fig. 2.**
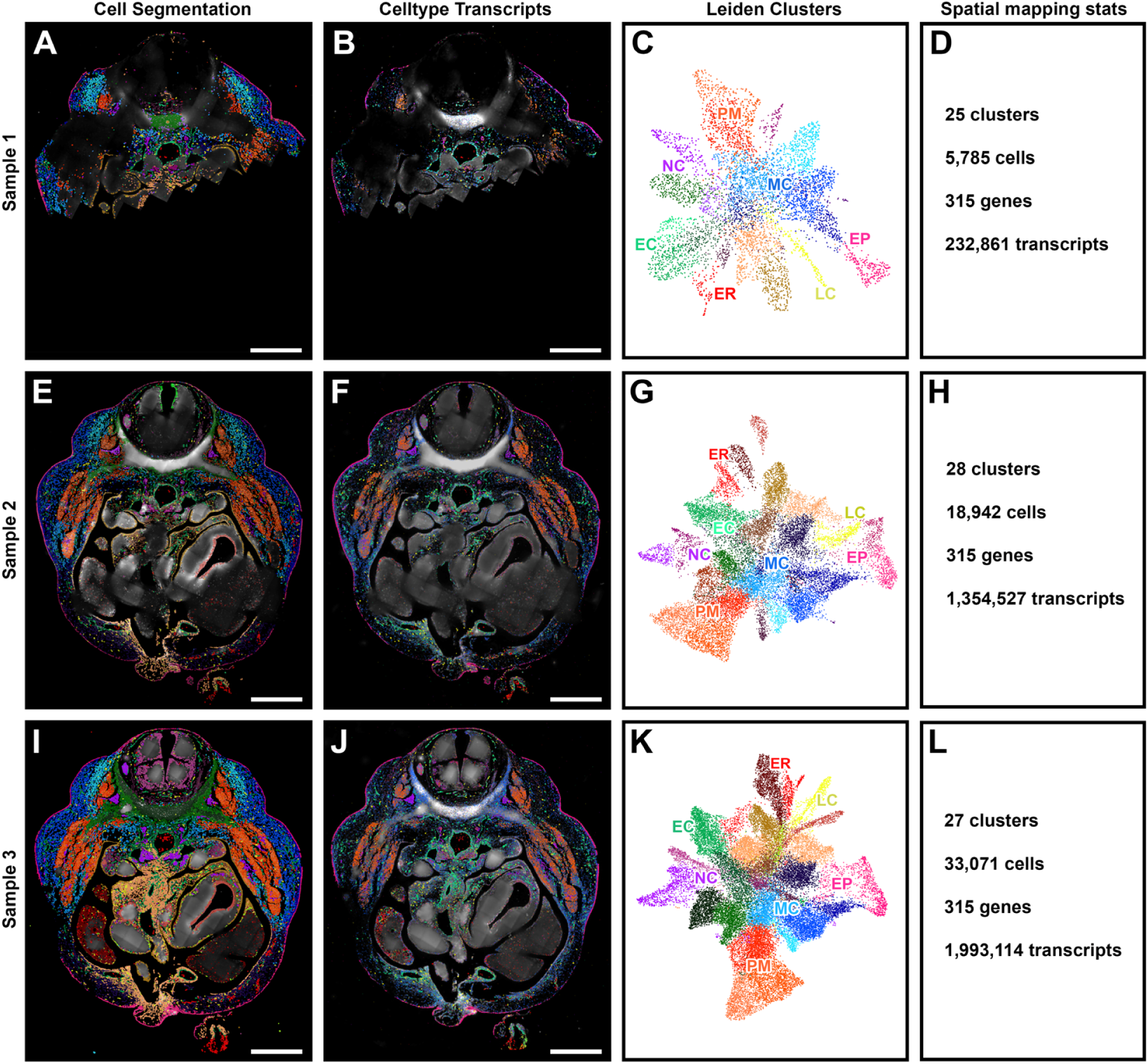
Wild-type spatial *in situ* imaging samples. (**A-L**) Three transverse E12 embryo sections (samples 1 – 3) processed on the MERSCOPE platform for spatial *in situ* imaging (285 genes, 30 controls). (**A,E,I**) Cell segmentation panels show Cellpose segmented cells, coloured according to correlation with sc-multiome and *de novo* clusters. (**B,F,J**) Celltype transcripts panel shows marker transcripts associated with cell types, coloured per cell type. (**C,G,K**) Spatial Leiden cluster UMAPs coloured accordingly and labelled by cell types. (**D,H,L**) Summary of spatial mapping stats per sample. Scalebars 500 µm. MC mesenchymal cells, PM paraxial mesoderm, NC neural crest derived, EC endothelial cells, EP epithelia, LC lymphocytes, ER erythrocytes.

**Extended Data Fig. 3.**
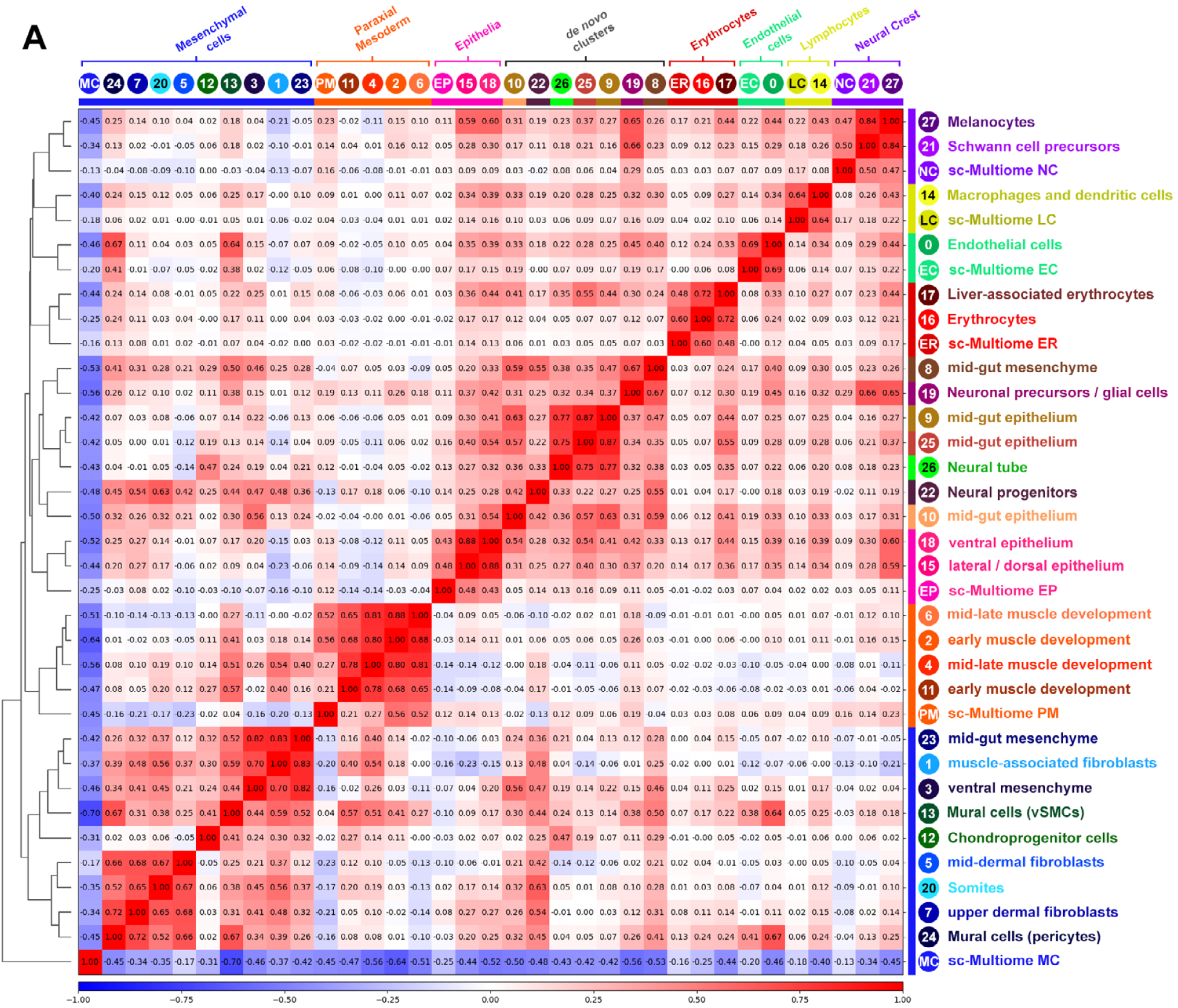
Correlation of marker genes between sc-multiome and spatial *in situ* imaging (wild-type sample 2). (**A**) Pairwise correlation matrix for sc-multiome (7) celltypes and spatial Leiden (28) clusters, using marker genes for each cell type. Spatial clusters derived from tissue outside the region profiled for sc-multiome are labelled as *de novo* clusters. Marker gene correlation together with spatial localisation enabled descriptive annotations of spatial clusters.

**Extended Data Fig. 4.**
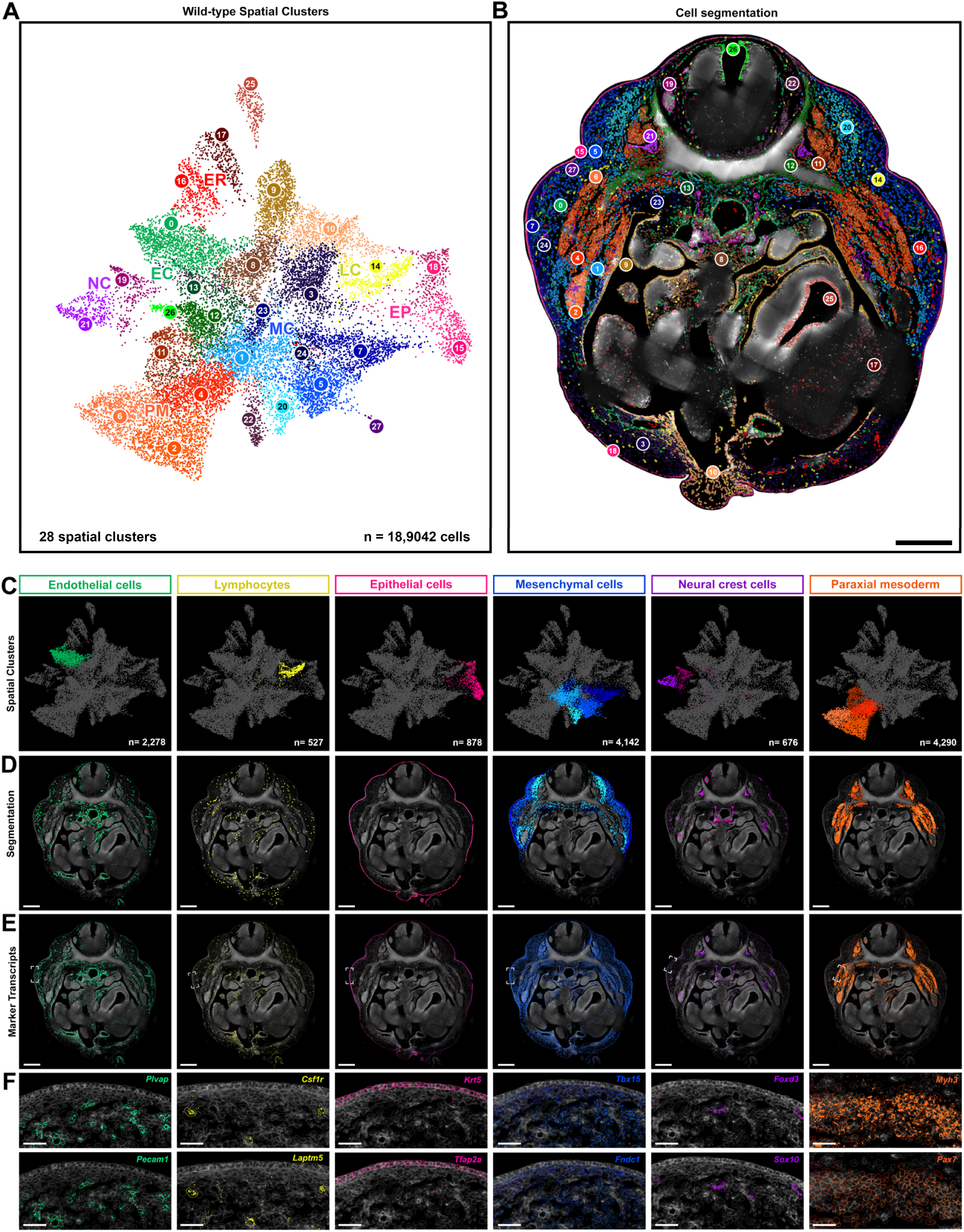
Spatial *in situ* imaging of cell types during early skin development in wild-type embryos (sample 2). (**A,B**) Segmented cells displayed as a UMAP and spatial *in situ* for wild-type sample 2, coloured according to Leiden clusters and cell types. (**C-F**) Summary of spatial mapping, showing cell type segmentation UMAPs (**C**), spatial imaging of cell types (**D**), and spatial expression of marker transcripts per cell type (**E**). (**F**) Expression of individual cell type marker transcripts in relation to segmented cell types. Scalebars: B,D,E, 200µm; F, 50µm.

**Extended Data Fig. 5.**
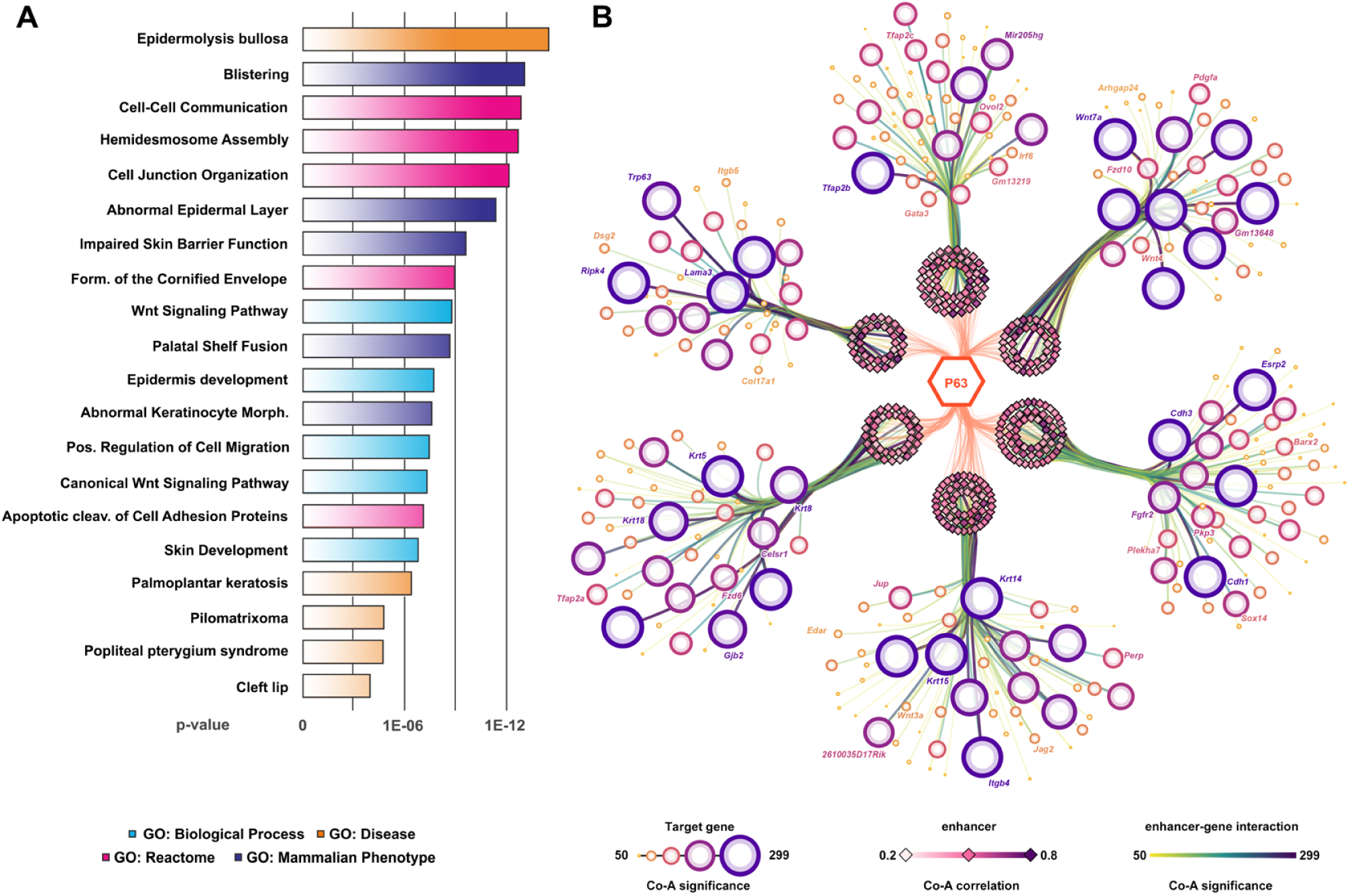
P63 predicted enhancer-gene regulatory network and gene ontology. (**A,B**) P63 predicted enhancer-gene regulatory network (eGRN) linking 4,909 enhancers with a P63 motif to 2,485 target genes. (**A**) Gene ontology terms of most significant predicted target genes (significance > 20; 1528 enhancers to 854 genes). (**B**) P63 directed network graph showing subset of most significant epithelial targets. P63 *cis*-regulatory enhancers (diamonds) are linked to gene promoters (circles), grouped by chromosome.

**Extended Data Fig. 6.**
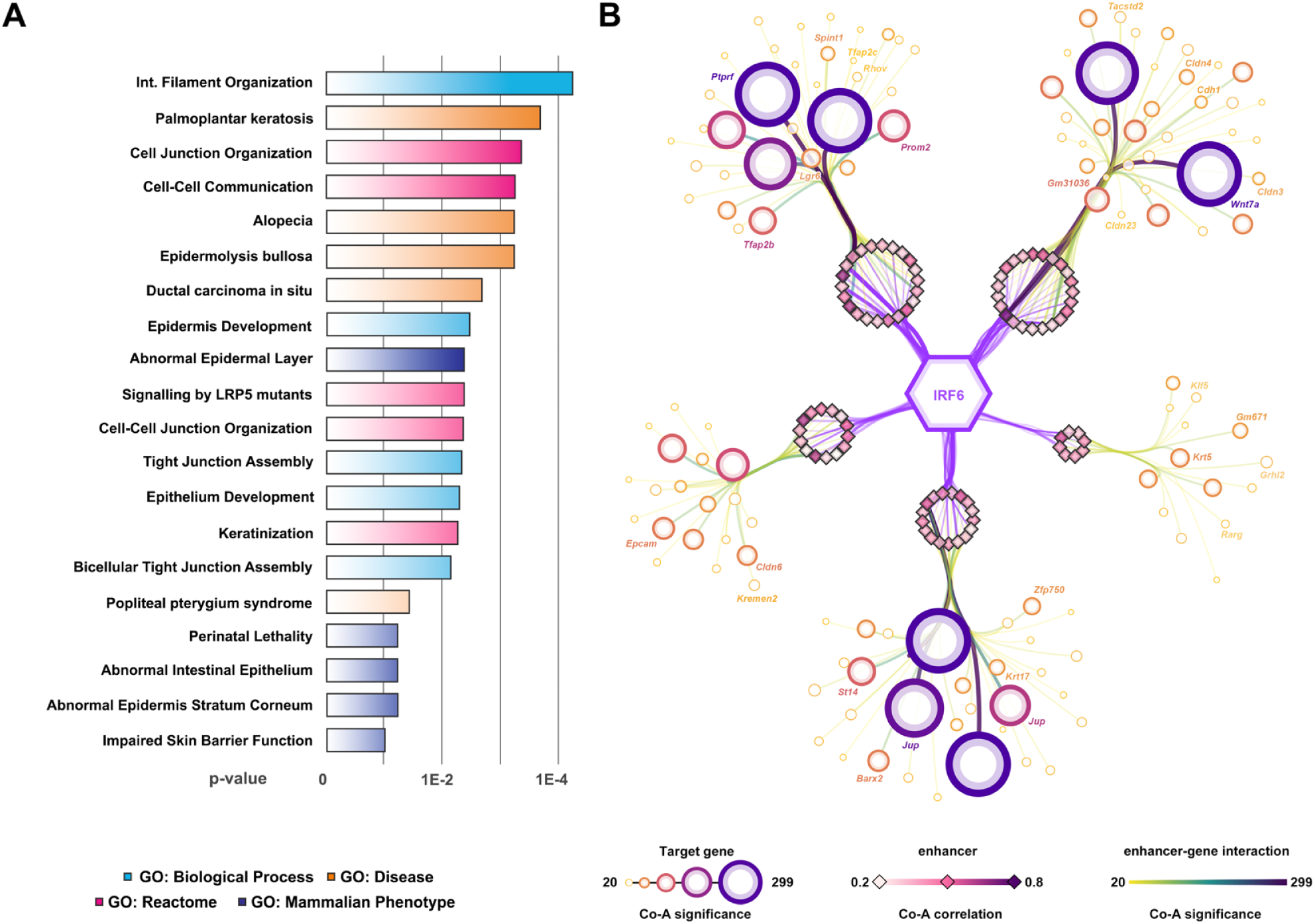
IRF6 predicted enhancer-gene regulatory network. (**A,B**) IRF6 predicted enhancer-gene regulatory network (eGRN) linking 722 enhancers with an Irf6 motif to 625 target genes. (**A**) Gene ontology terms of predicted target genes. (**B**) IRF6 directed network graph showing a subset of most significant epithelial targets. IRF6 *cis*-regulatory enhancers (diamonds) are linked to gene promoters (circles).

**Extended Data Fig. 7.**
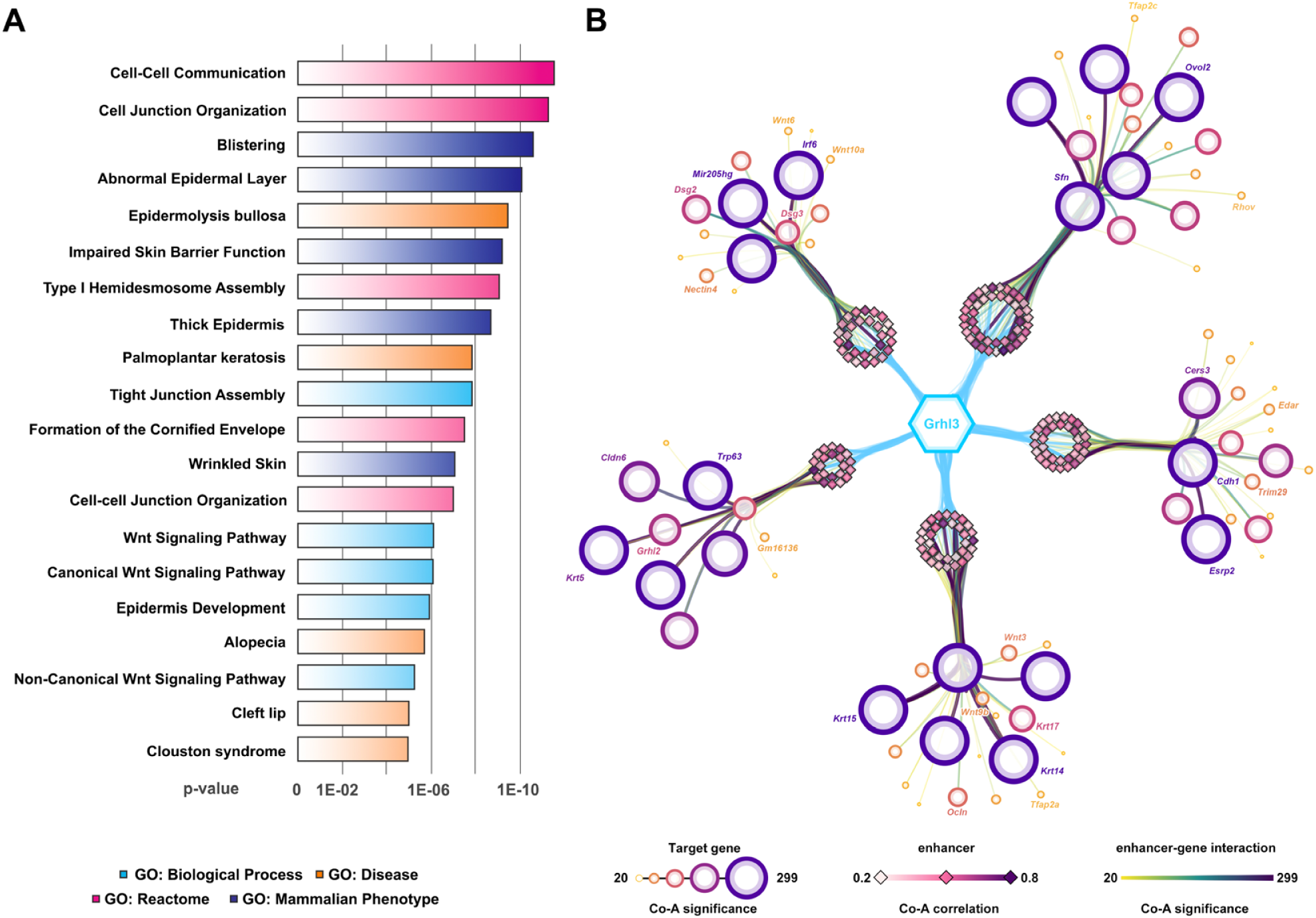
GRHL3 predicted enhancer-gene regulatory network. (**A,B**) GRHL3 predicted enhancer-gene regulatory network (eGRN) linking 3,971 enhancers with a GRHL3 motif to 2,337 target genes. (**A**) Gene ontology terms of most significant predicted target genes (significance > 20; 1193 enhancers to 786 genes). (**B**) GRHL3 directed network graph showing subset of most significant epithelial targets. GRHL3 *cis*-regulatory enhancers (diamonds) are linked to gene promoters (circles).

**Extended Data Fig. 8.**
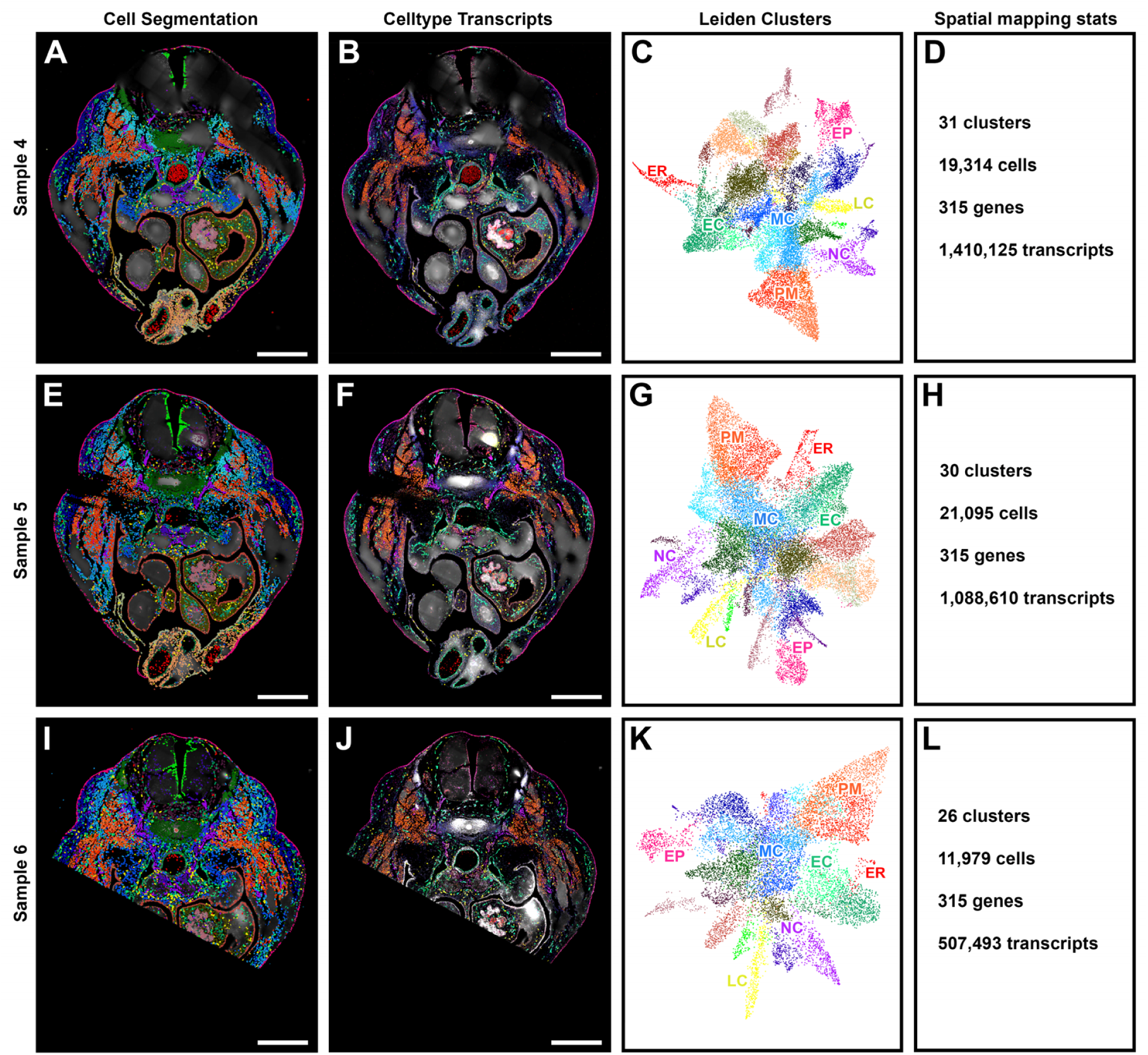
*Trp63*^-/-^ spatial *in situ* imaging samples. (**A-L**) Three transverse E12 embryo sections (samples 4 - 6) processed on the MERSCOPE platform for spatial *in situ* imaging (285 genes, 30 controls). (**A,E,I**) Cell segmentation panels show Cellpose segmented cells, coloured according to correlation with sc-multiome and *de novo* clusters. (**B,F,J**) Celltype transcripts panel shows marker transcripts associated with cell types, coloured per cell type. (**C,G,K**) Spatial Leiden cluster UMAPs coloured accordingly and labelled by cell types. (**D,H,L**) Summary of spatial mapping stats per sample. Scalebars 500µm. MC mesenchymal cells, PM paraxial mesoderm, NC neural crest derived, EC endothelial cells, EP epithelia, LC lymphocytes, ER erythrocytes.

**Extended Data Fig. 9.**
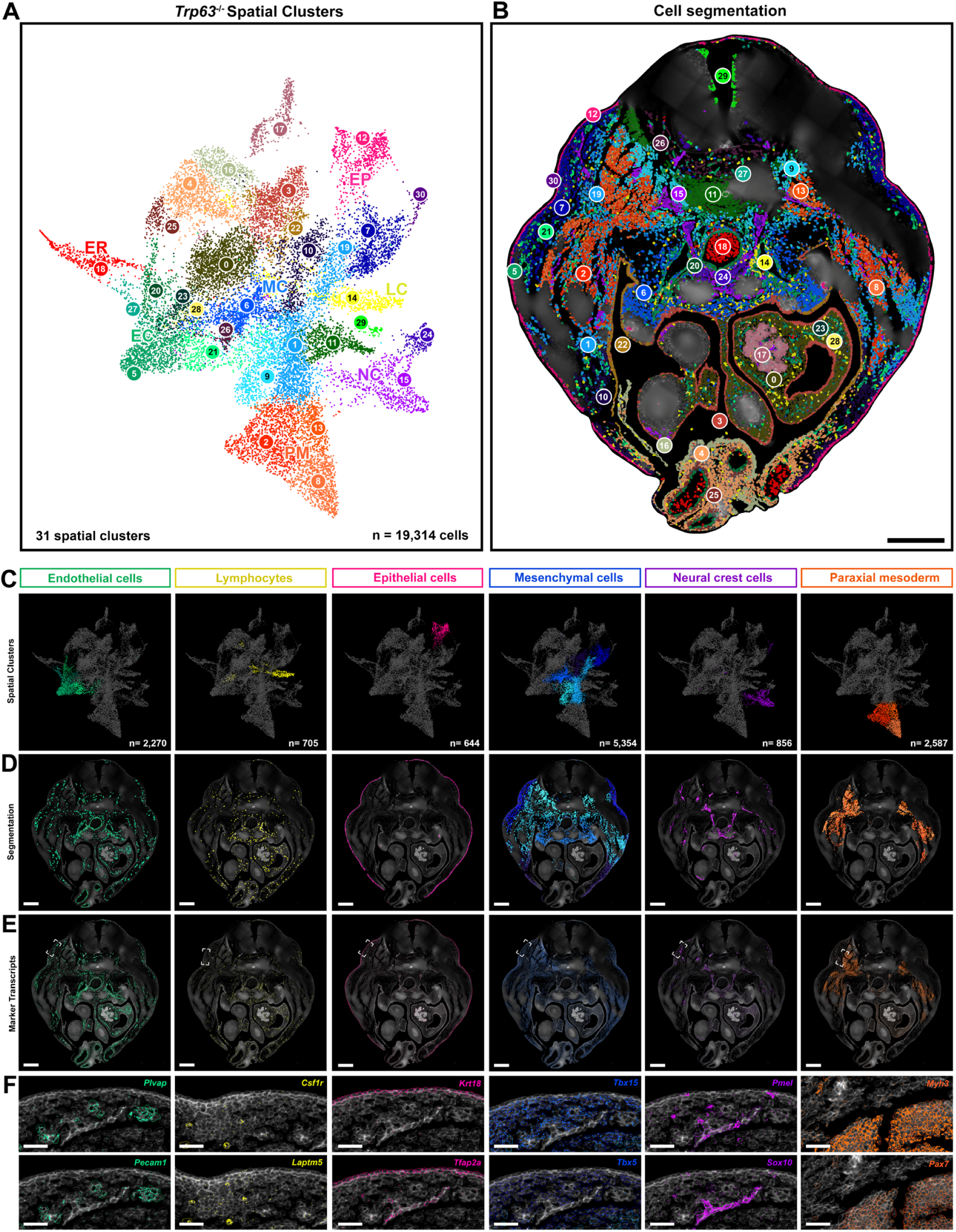
Spatial *in situ* imaging of cell types during early skin development in *Tp63*^-/-^ embryos (sample 4). (**A,B**) Segmented cells displayed as a UMAP and spatial *in situ* for *Tp63*^-/-^ sample 4, coloured according to Leiden clusters and cell types. (**C-F**) Summary of spatial mapping, showing cell type segmentation UMAPs (**C**), spatial imaging of cell types (**D**), and spatial expression of marker transcripts per cell type (**E**). (**F**) Expression of individual cell type marker transcripts in relation to segmented cell types. Scalebars: B,D,E, 200µm; F, 50µm.

**Extended Data Fig. 10.**
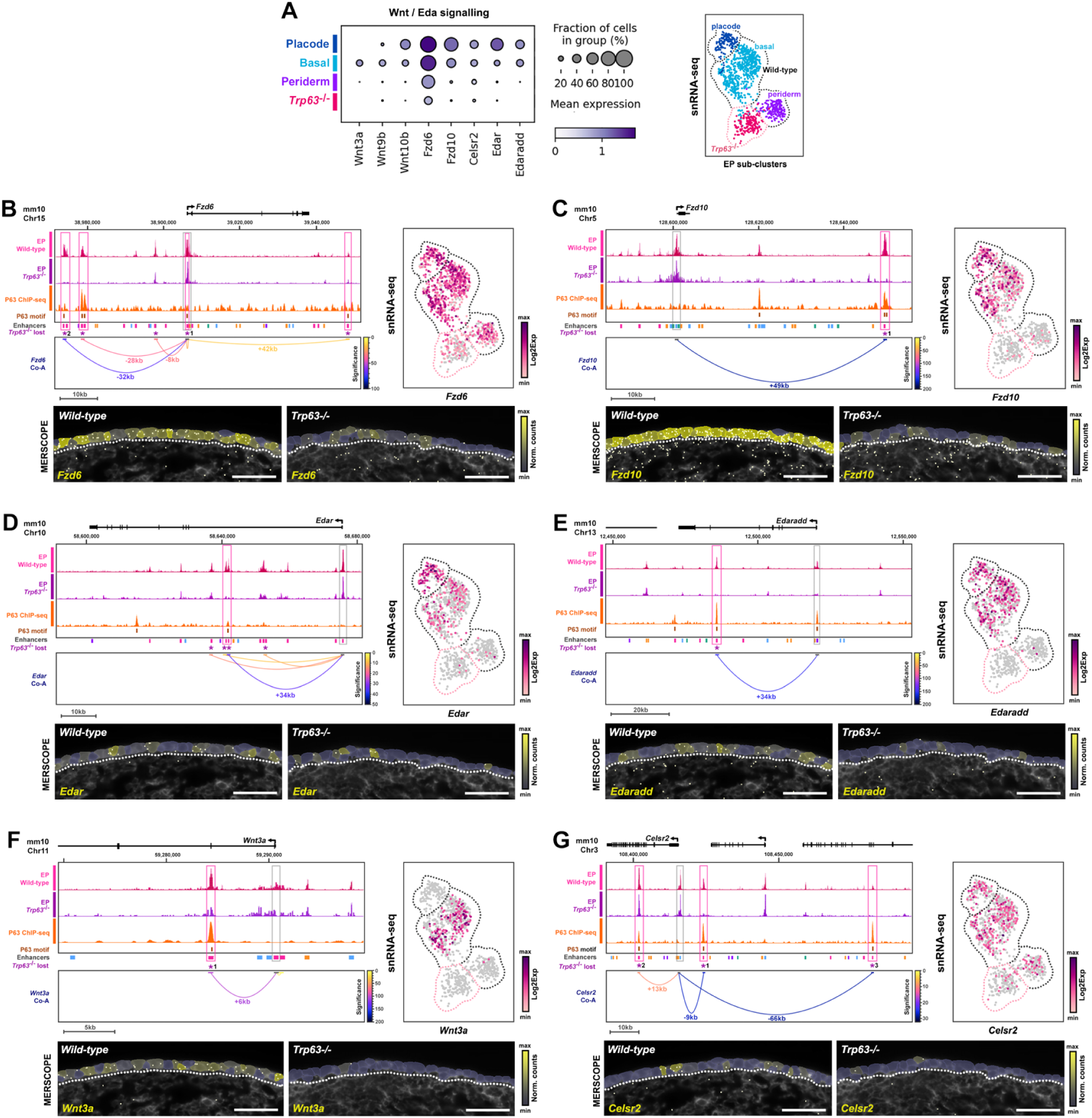
P63 regulates Wnt and Edar signalling pathway components. (**A**) sc-multiome gene expression dot plots of predicted P63 targets, associated with Wnt and Edar signalling. (**B-G**) UCSC genome (mm10) screenshots of *Fzd6* (**C**), *Fzd10* (**D**), *Edar* (**E**), *Edaradd* (**F**), *Wnt3a* (**G**), and *Celsr2* (**H**) showing epithelial (EP) snATAC-seq signals for wild-type (pink) and *Trp63*^-/-^ (purple) samples, P63 ChIP-seq (orange), co-accessible (Co-A) interactions and comparison of snRNA-seq and spatial expression of target gene transcripts. Asterisk (*) denotes lost interaction in *Trp63*-/- multiome. Scale bars 50µm

**Extended Data Fig. 11.**
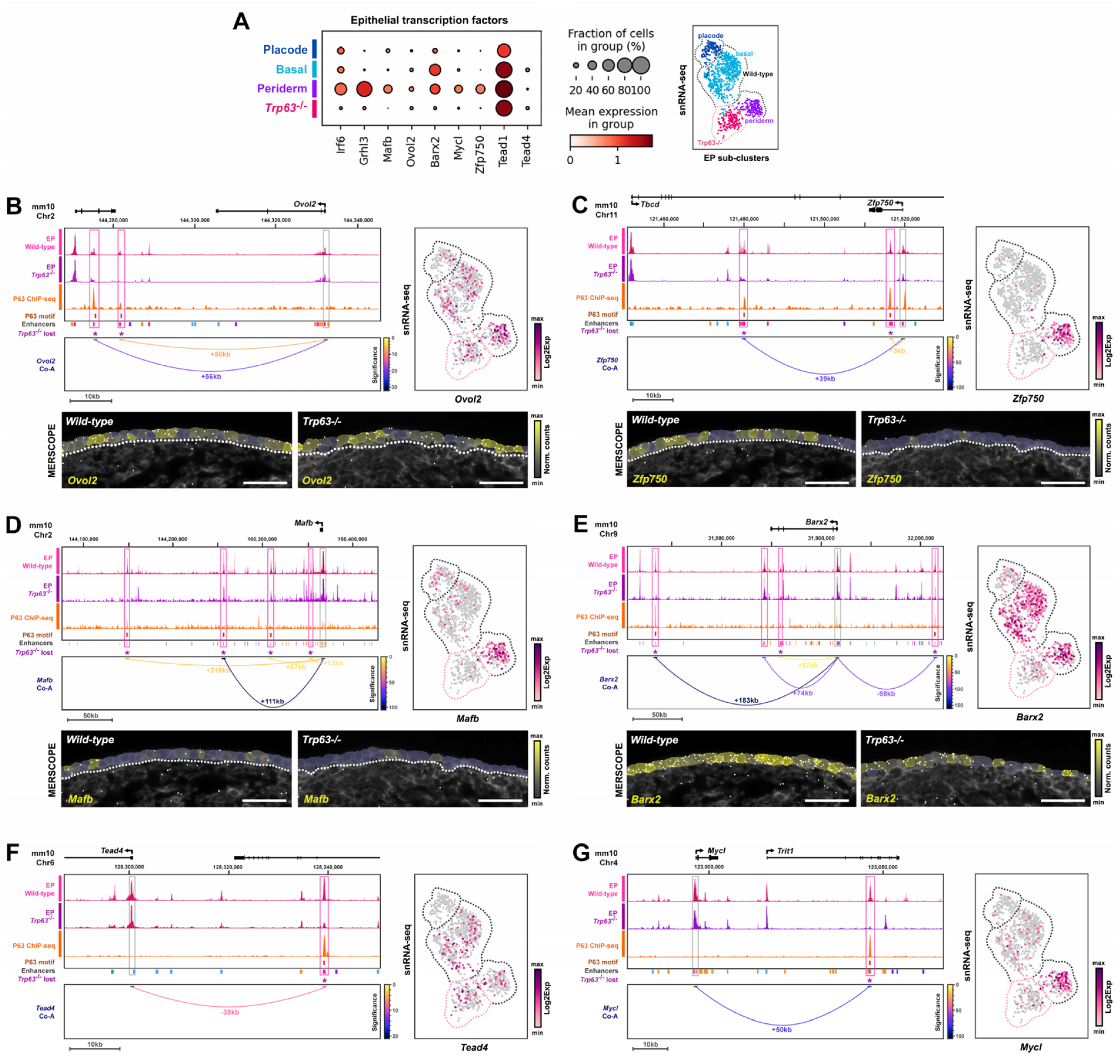
P63 regulates an epithelial transcription factor network. (**A**) sc-multiome gene expression dot plots of predicted P63 transcription factor targets. (**B-G**) UCSC genome (mm10) screenshots of *Ovol2* (**B**), *Zfp750* (**C**), *Mafb* (**D**), *Barx2* (**E**), *Tead4* (**F**) and *Mycl* (**G**) showing epithelial (EP) snATAC-seq signals for wild-type (pink) and *Trp63*-/- (purple) samples, P63 ChIP-seq (orange), co-accessible (Co-A) interactions and comparison of snRNA-seq and spatial expression of target gene transcripts. Asterisk (*) denotes lost interaction in *Trp63*-/- multiome. Scale bars 50µm

**Extended Data Fig. 12.**
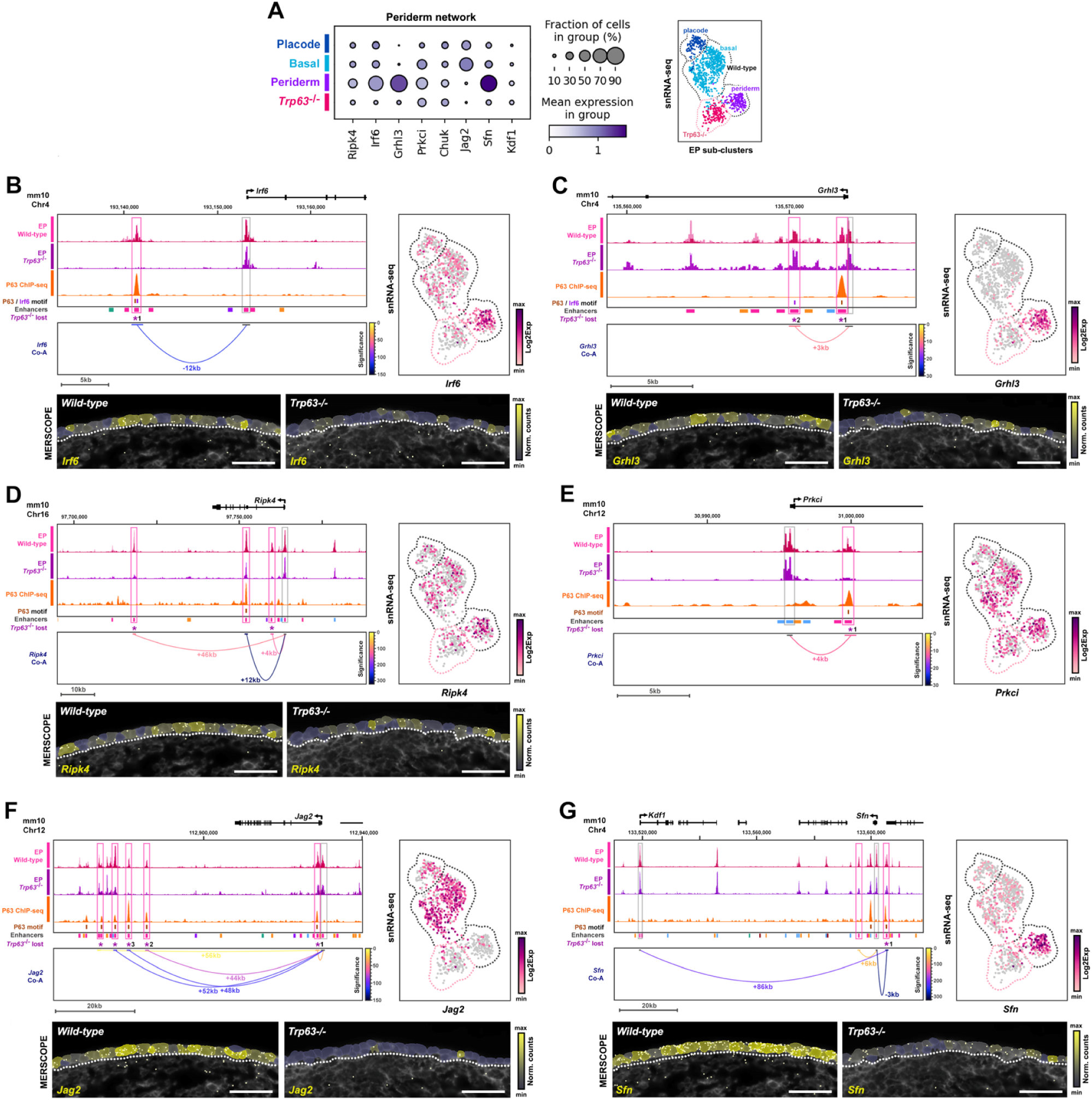
P63 regulates critical genes implicated in periderm gene regulatory network. (**A**) sc-multiome gene expression dot plots of predicted P63 targets associated with periderm gene regulatory network. (**B-G**) UCSC genome (mm10) screenshots of *Irf6* (**C**), *Grhl3* (**D**), *Ripk4* (**E**), *Prkci* (**F**), *Jag2* (**G**), and *Kdf1*/*Sfn* (**H**) showing epithelial (EP) snATAC-seq signals for wild-type (pink) and *Tp63*-/- (purple) samples, P63 ChIP-seq (orange), co-accessible (Co-A) interactions and comparison of snRNA-seq and spatial expression of target gene transcripts. Asterisk (*) denotes lost interaction in *Tp63*-/- multiome.

**Extended Data Fig. 13.**
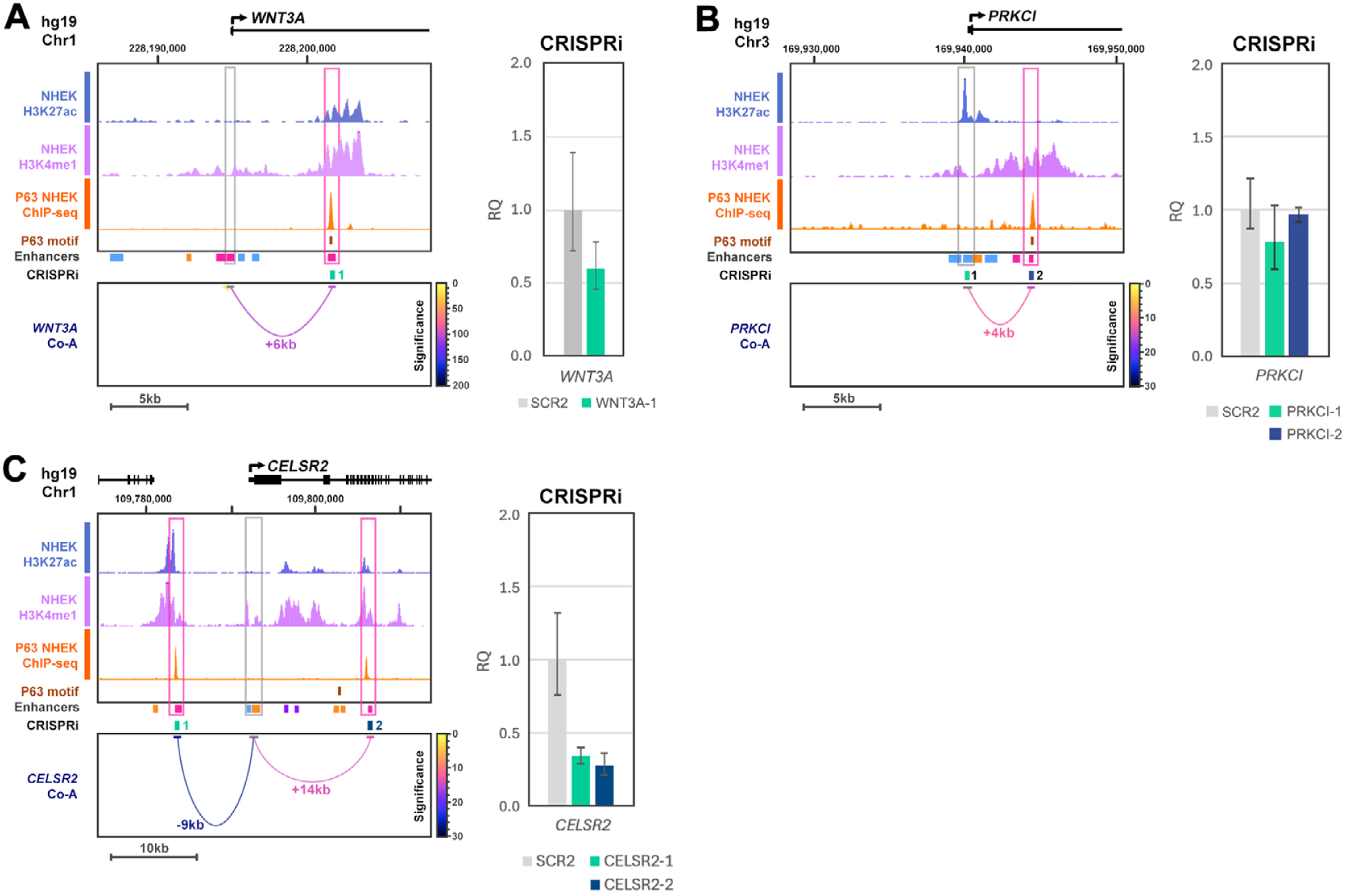
Comparative functional analyses of P63 enhancer-gene targets. (**A-C**) UCSC genome (hg19) screenshots of *WNT3A* (**A**), *PRKCI* (**B**) and *CELSR2* (**C**) showing conserved *cis*-regulatory interactions associated with histone methylation profiles (ChIP-seq; blue, purple) and P63 occupancy (ChIP-seq; orange) in epithelial cell lines. For each locus, the indicated enhancer was targeted with gRNA for CRISPRi, with gene expression (RQ) quantified (qPCR) relative to a scrambled control (SCR2). Each bar represents a single transfected cell line. Error bars denote RQ expression min/max confidence intervals (95%).

